# DNA sequence context and the chromatin landscape differentiate sequence-specific transcription factor binding in the human malaria parasite, *Plasmodium falciparum*

**DOI:** 10.1101/2023.03.31.535174

**Authors:** Victoria A. Bonnell, Yuning Zhang, Alan S. Brown, John Horton, Gabrielle A. Josling, Tsu-Pei Chiu, Remo Rohs, Shaun Mahony, Raluca Gordân, Manuel Llinás

## Abstract

Development of the human malaria parasite, *Plasmodium falciparum,* is regulated by a limited number of sequence-specific transcription factors (TFs). However, the mechanisms by which these TFs recognize genome-wide binding sites is still largely unknown. To address TF specificity, we investigated the binding of two TF subsets that either bind CACACA or GTGCAC DNA sequence motifs and further characterized PfAP2-G and PfAP2-EXP which bind unique DNA motifs (GTAC and TGCATGCA). We interrogated the impact of DNA sequence and chromatin context on *P. falciparum* TF binding by integrating high-throughput *in vitro* and *in vivo* binding assays, DNA shape predictions, epigenetic post-translational modifications, and chromatin accessibility. We determined that DNA sequence context minimally impacts binding site selection for CACACA-binding TFs, while chromatin accessibility, epigenetic patterns, co-factor recruitment, and dimerization contribute to differential binding. In contrast, GTGCAC-binding TFs prefer different DNA sequence context in addition to chromatin dynamics. Finally, we find that TFs that preferentially bind divergent DNA motifs may bind overlapping genomic regions *in vivo* due to low-affinity binding to other sequence motifs. Our results demonstrate that TF binding site selection relies on a combination of DNA sequence and chromatin features, thereby contributing to the complexity of *P. falciparum* gene regulatory mechanisms.

**Key Points:**

- Sequence and chromatin context determine differential DNA-binding specificity of *P. falciparum* TFs
- TFs with paralogous DNA-binding domains in *P. falciparum* are not functionally redundant
- TFs with differing sequence-specificity can co-occupy sites through low-affinity DNA interactions

## Introduction

Sequence-specific transcription factors (TFs) bind a core DNA sequence motif through base-specific contacts using the major and/or minor groove of DNA^1–6^. However, the nature of this protein-DNA interaction is more complex than simply recognizing a specific DNA motif, since for any given TF only a fraction of the total possible genome-wide sites are bound^1, 7–9^. Additional features such as DNA sequence context, local DNA topography, post-translational modifications (PTMs) of the TF or histones, TF protein concentration, TF timing of expression, DNA methylation patterns, protein-interaction partners, and chromatin state can all influence TF binding site recognition^1, 5, 10–17^. Determining how individual TFs select and bind to cognate DNA motifs *in vivo* is central to understanding gene regulatory networks, since dysregulation of TF binding can be deleterious as seen in diseases such as cancer (*i.e.,* P53 and MYC)^18, 19^ and can impact stress tolerance in plant crops (*i.e.,* AP2/ERFs)^20–23^.

Most eukaryotes encode expanded TF families that arise through gene duplication events and subsequent diversification, resulting in evolutionarily conserved DNA-binding domains (DBDs) that bind highly similar DNA sequences (*i.e.,* paralogous domains)^1, 7, 8, 24, 25^. Paralogous TFs are highly represented in model eukaryotes from unicellular (*e.g.,* yeast) to multicellular (*e.g.,* mammals) organisms, have been shown to function in both unique or redundant manners, and govern alternate transcriptional regulatory networks in different cell types^25–27^. The most well-studied example is the homeobox domain (HOX) TF family in animals, which contains many paralogous TFs that all recognize A/T-rich DNA motifs^28^. While HOX TFs recognize similar DNA motifs *in vitro*, they have divergent *in vivo* functions. This is largely due to moderate-and low-affinity binding driven by a combination of factors including the DNA sequence context surrounding the A/T-rich HOX motifs, interactions with divergent co-factors inducing latent specificities, and varying abilities to bind inaccessible chromatin^11, 17, 28–30^.

Single-celled eukaryotes, such as yeast, have served as models to explore the unique expansion of paralogous TF binding in the absence of multicellularity^31–36^. Yeast paralogous TFs generally regulate vastly differing target genes, often in response to diverse extracellular environments^37, 38^. Similarly, single-celled, eukaryotic Apicomplexan parasites encode TFs with paralogous DBDs. However, the number of paralogous domains are drastically reduced due to genome reduction following evolutionary adaptation to a parasitic lifestyle^39, 40^. The Apicomplexan human malaria parasite, *Plasmodium falciparum,* presents a unique opportunity to explore the challenge of paralogous protein-DNA specificity in the context of a 22.9Mb genome that is one of the most A/T-rich genomes sequenced to date (ranging from 85% genome-wide up to 90% A/T in intergenic regions)^41^. Despite this, *P. falciparum* has evolved surprisingly few TFs that bind A/T-rich motifs and possesses a limited repertoire of sequence-specific TFs^42, 43^. These TFs are thought to act through an array of unique DNA sequence motifs found in regulatory regions upstream of transcription start sites (TSSs)^41–47^.

The *P. falciparum* genome encodes homeodomain-like (HD)^48^, myeloblastosis (MYB)^49, 50^, high mobility group box (HMGB)^51, 52^ TFs as well as an expanded set of diverse zinc finger domain proteins^53–55^ that have not been extensively characterized to date^41, 45, 47^. However, the largest defined family of *P. falciparum* TFs (28 members) is the Apicomplexan APETALA2 (ApiAP2) family of DNA-binding proteins that contain one to three AP2 DBDs^42, 43, 56–58^. APETALA2/Ethylene response factor (AP2/ERF) TFs are also one of the most highly represented TF families in plant-lineage genomes. AP2/ERF protein-DNA recognition occurs via three anti-parallel beta-strands directly interacting with the DNA major groove, a recognition modality that is consistent in both plant and Apicomplexan AP2/ERF protein-DNA complexes^59–65^. Plant AP2/ERF TFs have highly paralogous DBDs that all recognize a GCC-box DNA motif, with well-defined activation and repression domains^23, 59–61^. In contrast, ApiAP2 proteins have more divergent AP2 domains that recognize a wide variety of unique DNA motifs and contain surprisingly few additional and discernable functional domains^43, 66^. *Plasmodium* ApiAP2 proteins have been identified as critical transcriptional regulators of virtually all developmental processes of the malaria parasite lifecycle^57, 58^, yet little is known about how ApiAP2 genome-wide DNA-binding selectivity is established.

Previous work has defined DNA sequence motifs recognized by most of the *P. falciparum* AP2 DBDs^42, 43^. These previous experiments did not account for either the effects of the A/T-rich *P. falciparum* sequence context or chromatin-based contributors. Therefore, in this study, we interrogated the relevance of both sequence and chromatin context on the genomic binding site selection of several paralogous and non-paralogous DBDs from TFs in *P. falciparum*. We generated a novel *P. falciparum*-specific genomic-context protein-binding microarray (gcPBM) to simultaneously probe protein-DNA interactions between DBDs and all intergenic instance of its cognate DNA motif directly from the *P. falciparum* genome^10, 12^. We also used comparative bioinformatic analyses to explore features of the chromatin environment such as genome-wide TF occupancy, chromatin accessibility, and epigenetic histone PTMs using new and published *P. falciparum* genome and epigenome datasets^44, 67–71^. Our findings suggest that a subset of *P. falciparum* TFs are greatly impacted by DNA sequence context while others are influenced by a complex interplay between TF timing of expression, genome-wide occupancy, and the chromatin landscape in a stage-dependent manner during parasite development.

*P. falciparum* critically relies on precise regulation of gene expression throughout its lifecycle^72–75^. In addition to the 48-hour asexual replicative cycle in human erythrocytes, *P. falciparum* parasites undergo several major developmental transitions, including sexual development for transmission between human and mosquito as well as growth and replication in human hepatocytes, the mosquito midgut, and mosquito salivary glands. These transformations all rely on the action of sequence-specific TFs in concert with epigenetic regulation^75, 76^ to program cellular differentiation, genome maintenance, immune system evasion, and development of the parasite^57, 58^. Therefore, our findings have implications for understanding the regulatory role of TFs critical to parasite development that may also serve as potential therapeutic targets.

## Methods

### Protein induction and affinity purification

All experiments were conducted using purified AP2 DNA-binding domains (DBDs) (starred [*] in **Figure 1A**; **Supplemental Figure 1A**) fused to an N-terminal glutathione-S-transferase (GST) tag and was purified as demonstrated previously^42, 43^. Briefly, each AP2 domain was previously cloned into the pGEX-4-T1 vector (GE Life Sciences) and transformed into BL21- CodonPlus(DE3)-RIL *E. coli* (Stratagene) for protein production using Isopropyl β-d-1- thiogalactopyranoside (IPTG) induction^43^. Verification of a successful GST-tagged AP2 protein purification by GST-affinity purification (Thermo Scientific Pierce Glutathione Superflow Agarose beads) was demonstrated by running protein samples on 4-15% stacking SDS-PAGE gel and stained with Coomassie Blue. Integrity of the GST tag was subsequently checked via western blotting of the purified recombinant protein using anti-GST antibodies (Invitrogen 71-7500) [1:1,500 dilution]. The homeodomain from HDP1 was tagged and purified similarly with modifications^48^ and gifted to us for the gcPBM experiments from the Kafsack lab at Cornell University.

**Figure 1:**
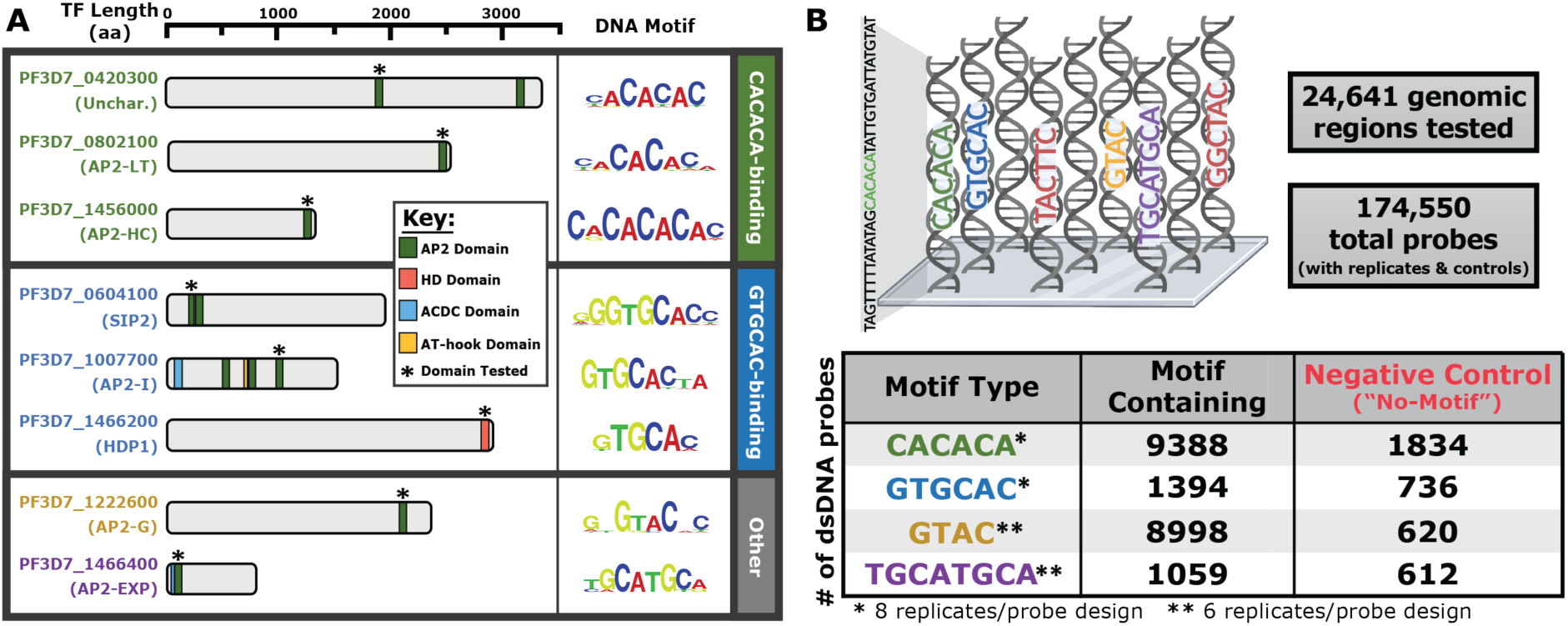
Multiple *P. falciparum* TFs with overlapping sequence preferences and design of the *P. falciparum* genomic-context protein-binding microarray (gcPBM) **(A)** Graphical representation of each TF examined in this study categorized into “CACACA-binding”, “GTGCAC-binding”, and “Other”. Protein lengths (in number of amino acids) are drawn to scale. Predicted protein domains were determined using NCBI Conserved Domain Search^90^ or defined by previous literature^43, 48^. Position weight matrix (PWM) logos are from previously published universal protein-binding microarray (PBM) experiments^42, 43, 48^. * Denotes the DNA-binding domains (DBDs) tested in this study; **(B)** Graphical representation of the *P. falciparum* gcPBM design. Position weight matrix (PWM) data^42, 43, 48^ was searched against intergenic regions of the *P. falciparum* genome (Pfalciparum3D7; version 3, release 38)^79^ and categorized into four motif types (CACACA, GTGCAC, GTAC, and TGCATGCA). All sequences were replicated eight(*) or six(**) times. Microarray graphic generated using BioRender.

### Universal Protein-Binding Microarray (PBM) for quality control

Each purified GST-tagged DBD stock was validated for DNA-binding specificity using 4×44k universal protein-binding microarrays (PBMs) (Agilent Technologies), as described previously^42, 43, 48, 77, 78^, before testing on the *Plasmodium falciparum*-specific genomic-context protein-binding microarray (gcPBM) design. The experiments were repeated, as previously demonstrated^42, 43^, with the exception that they were tested on protease-treated, re-used arrays, to determine the amount of protein necessary to generate signal within the linear range of the GenePix 4300B Microarray Scanner (Molecular Devices) at a resolution of 5µm. In brief, the microarray stripping protocol^77^ includes: (1) an overnight incubation in a protease solution (350 units of protease [Sigma 537088], 10%v/v SDS, and 10mM EDTA), 3 washes in 1xPBS with 0.5%v/v Tween20, a final rinse in 1xPBS, then scanning for Alexa488 signal to ensure digestion of protein from the previous experiment. The PBM method^77^ includes 5 major steps with 1xPBS with 0.01- 0.5%v/v Tween20 washes in between each step: (1) Blocking the microarray with 2%w/v milk in 1xPBS for one hour in the dark, (2) Applying the protein-binding mixtures to the microarray for one hour, (3) Probing protein-DNA interactions with an anti-GST, rabbit IgG, Alexa488- conjugated antibody (1:40 dilution in 2%w/v milk blocking solution; Invitrogen, A11131) for one hour, (4) Scanning the microarray for Alexa488 signal (GenePix Pro version 7.2 software), and (5) Identifying enriched DNA motif using a gapped 8-mer DNA motif-searching perl script modified from original script^77^ to normalize only based on local neighboring probes instead of Cy3 signal introduced from double-stranding the microarray.

### Design of Plasmodium falciparum Genomic-Context Protein-Binding Microarray (gcPBM)

Using position weight matrix (PWM) data from published work^42, 43^, all instances of each motif were identified in the *P. falciparum* genome (*Plasmodium falciparum* 3D7 strain genome release v38^79^), using a motif E-score cutoff of >0.45^10, 80^. Only intergenic regions (excluding telomeric regions) were used for this gcPBM design. The numbers of probes found per DNA motif are as follows: 2848 probes with putative sites for PF3D7_0420300_D1 (500 negative controls), 4251 probes with putative sites for AP2-LT (500 negative controls), 3864 probes with putative sites for PF3D7_1305200 (500 negative controls), 4321 probes with putative sites for AP2-HC (500 negative controls), 1459 probes with putative sites for SIP2_D1 (500 negative controls), 3742 probes with putative sites for AP2-I_D3 (500 negative controls), 8998 probes with putative sites for AP2-G (1000 negative controls), and 1059 probes with putative sites for AP2-EXP (1000 negative controls). HDP1 motif-specific genomic sequences were not initially included in the gcPBM due to its identification and characterization^48^ after the initial design of the gcPBM experiment, but was included due to PWM similarities to SIP2_D1 and AP2-I_D3. Any sequence containing another instance of the centered motif in the left or right flanks was mutated to prevent multiple binding sites per 36-bp window. Due to the similarities between the CACACA and GTGCAC PWMs, there were genomic DNA sequences that led to redundant probe designs, which were discarded, leaving only one instance of the sequence. After discarding redundant probe designs with motif types, the total number of probes per motif type was as follows: 9388 probes with putative CACACA sites (1834 CACACA negative controls); 1394 probes with putative GTGCAC sites (736 GTGCAC negative controls); 8998 probes with putative GTAC sites (620 GTAC negative controls); and 1059 probes with putative TGCATGCA sites (612 TGCATGCA negative controls). Overall, the *P. falciparum* gcPBM design reached a total of 24,641 unique genomic regions (**Supplemental File 1**). Each double stranded DNA probe was represented in both the 5′ and 3′ orientations, with one end of each DNA molecule attached to a glass slide (**Supplemental Figure 2A**). Additionally, each DNA probe was replicated in random areas of the microarray surface (four CACACA/GTGCAC replicates and three GTAC/TGCATGCA replicates per orientation), which brought the total number of DNA probes to 174,550 spots for a 4×180k microarray (Agilent Technologies). Additional spots on the array were set aside for control grid alignment, microarray scanning, and downstream analysis.

### Genomic-Context Protein-Binding Microarray (gcPBM) experiment

The single-stranded DNA microarrays were double-stranded by solid-state primer extension as reported previously^10, 12^. The 24bp primer sequence (5‵-GTCTTGATTCGCTTGACGCTGCTG- 3‵) was used to double-strand all gcPBM slides for this study. Each GST-tagged DBD was tested for DNA sequence specificity by applying protein to the *P. falciparum* gcPBM as demonstrated previously^10, 77, 80^. The major components of the protein-binding mixture are as follows: 1%w/v milk, 0.2mg/mL BSA, 0.5%v/v salmon testes DNA, and 0.03%v/v TritonX-100 in 1xPBS^10, 12, 13, 77, 78^. Amounts of recombinant protein necessary for DNA-binding was empirically determined during preliminary universal PBM experiments detailed above (average amount added is around 25µg; identified by A280 signal on NanoDrop 2000 Spectrophotometer [Thermo Scientific]).

### Genomic-context Protein-Binding Microarray (gcPBM) data acquisition and analysis

Immediately after completing the gcPBM experiment, each microarray chamber was scanned using the GenePix 4400A Microarray Scanner (Molecular Devices) at a resolution of 2.5µm, with the 488nm wavelength laser, along with the GenePix Pro7 software to generate image files (.tif). The image files were aligned with the GenePix Array List (.gal) file to associate raw signal intensity to the *P. falciparum* genome-derived DNA sequences, which generates a GenePix Results (.gpr) file. The .gpr files were then further processed and normalized using the previously published Masliner script and downstream analysis made by the Bulyk lab and modified by the Gordân lab^10, 77, 78, 80^. Final values used for this study are a single data point representing the highest natural log median binding intensity value across the replicates between either the 5‵ or 3‵ orientations for each DNA sequence.

### Validation with Electrophoretic Mobility Shift Assays (EMSAs)

All gel shift experiments were conducted using the LightShift Chemiluminescent EMSA kit and protocol (Thermo Scientific), with some modifications. DNA sequences designed for each gel shift are found in **Supplemental File 2**. Single-stranded biotinylated DNA oligos were annealed to their reverse complement sequence to generate double-stranded DNA probes using the Duplex Buffer recipe (100mM Potassium acetate, 30mM HEPES [pH 7.5]) and protocol from Integrated DNA Technologies (IDT) website. The minimal concentration of protein added to the binding reaction to produce a robust shift was empirically determined by titrating in varying amounts of protein with the wildtype DNA sequences (data not shown). The standard protein-binding mixture includes: 1x Binding buffer (100mM Tris-HCl [pH 7.5], 500mM KCl, 10mM DTT), 5mM MgCl_2_, 25ng/µL Poly dI·dC, 0.05%v/v NP-40, protein in 25%v/v glycerol, and biotinylated dsDNA probe. Briefly, the protocol includes: (1) Combining all components of the protein-binding mixture and incubating at room temperature for 20 minutes, (2) running samples on a pre-run 6% non-denaturing PAGE gel using 0.5xTBE, (3) Transfer DNA onto a nylon membrane using 0.5xTBE, (4) Crosslink the nylon membrane at 312nm for 15 minutes with a transilluminator, (5) Block the membrane for 15 minutes with shaking, (6) Detect the biotin-labeled DNA by added Stabilized Streptavidin-Horseradish Peroxidase Conjugate (1:300 dilution) for 15 minutes with shaking in blocking buffer, (7) Wash the membrane 4× with shaking in 1X wash buffer, (8) Equilibrate the membrane with shaking, and (9) Incubate with the Substrate Working Solution (1:1 Luminol Enhancer/Peroxide) without shaking. Final images taken on ChemiDoc XRS+ Molecular Imager (Bio-Rad). Exposure times determined by minimal exposure without oversaturation.

### Designing DNA oligos with predicted DNA shape mutations

The DNA oligos used to investigate the impact of predicted *in vitro* DNA shape on binding were generated by the mutation design tool of TFBSshape^81^. TFBSshape produces oligo sequences with mutations that minimize DNA sequence changes while generating dynamics changes to the predicted shape features according to the distance between their wild type and mutant. The sequence distance is determined by Levenshtein distance that sums the number of substitutions, deletions or insertions required to transform from a mutant to its wild type sequence. The predicted shape distance is calculated in Euclidean distance between two normalized shape feature vectors for a wild type and its mutant sequence. The normalized predicted shape features including helix twist (HelT), minor groove width (MGW), propeller twist (ProT), and roll are derived from DNAshapeR^82^. Three bps on the flanks of the fixed core ‘AGTGCATTA’ were subjected to mutation, as shown in lowercase in **Supplemental File 3**. The oligos with the maximum shape distance, with respect to the preserved sequence distance were selected. The sequence distance was preserved, and the predicted shape distance was calculated and sorted among all possible mutations.

### Generation of Plasmodium falciparum parasite line and culturing conditions

To generate the PfAP2-LT^HA^ tagged parasite line, a 700bp homology region with a 3xHA sequence left-flanked by the 3‵-end of AP2-LT coding sequence and right-flanked by the endogenous 3‵- untranslated region (UTR) of AP2-LT was cloned into the pDC2-U6A-hDHFR vector. This CRISPR single-plasmid design contained a *pfap2-lt* locus-targeted guide RNA sequence with BbsI restriction site, Cas9^HA^, AMP^R^, WR^R^, and NotI and SacI restriction sites flanking the homology region. The final plasmid was transformed into heat shock competent DH5α cells with 100µg/mL ampicillin (AMP). Then the plasmid was purified and ethanol precipitated overnight at -80°C. Wild type parasites (3D7 strain) were cultured under standard *P. falciparum* culturing conditions (5%O_2_, 7% CO_2_, 37°C, RPMI 1640 media with 0.5% AlbumaxII and hypoaxanthine). The purified plasmid DNA (100µg) was transfected by electroporating uninfected erythrocytes and adding trophozoite-stage parasites to later invade the erythrocytes preloaded with plasmid DNA. After parasite reinvasion, parasites were treated with 2.5nM of WR99210 for one week and switched to no drug standard parasite media. After limited dilution cloning to generate a clonal parasite line, genomic DNA was purified from the AP2-LT^HA^ parasite culture (Qiagen) and verified for integration by PCR (primer sequences in **Supplemental File 2**), Sanger sequencing, and whole genome sequencing via high-throughput Illumina sequencing.

### AP2-LT Chromatin immunoprecipitation followed by high-throughput sequencing (ChIP-seq)

The ChIP-seq experiment was carried out using a similar protocol as previous published work^83^. The ChIP-seq protocol had 5 major steps including: (1) Chemically crosslinking the all protein-protein and protein-chromatin interactions, (2) Isolation of parasite nuclei, (3) Parasite nuclei lysis with chromatin sonication, (4) Protein-chromatin complex immunoprecipitation, and (5) DNA purification. The crosslinking step included: (a) Chemical crosslinking suspended *Pf*AP2-LT^HA^ (or WT culture for negative control experiment) parasite culture (at least 10^8^ 36-45hpi schizont-stage parasites synchronized with 10%w/v Sorbitol more than one cycle prior) with 1%v/v formaldehyde for 10 minutes at 37°C, (b) Quench the crosslinking with 125mM Glycine on ice for 5 minutes, (c) Pellet the cells by centrifugation and lyse the red blood cells with 0.1%w/v Saponin in 1xPBS, and (d) Wash the cells by repeated centrifugation with 1xPBS to remove the red blood cell debris. The nuclei isolation step included: (a) Resuspend parasite pellet with 10^9^ schizonts/2mL Lysis Buffer (10mM HEPES [pH 7.9], 10mM KCl, 0.1mM EDTA [pH 8.0], 0.1mM EGTA [pH 8.0], 1mM DTT (added just before using), and 1x protease inhibitors) and incubate on ice for 30 minutes, (b) Add a final concentration of 0.25%v/v NP-40 and incubate for 1 minute, (c) Lyse parasite membranes with pre-chilled glass dounce homogenizer for 100 strokes per 10^9^ schizonts/2mL Lysis Buffer, and (d) Pellet the nuclei and freeze pellet at -80°C overnight. The sonication step included: (a) Resuspend parasite nuclei pellet in 500µL of Shearing Buffer (0.1%v/v SDS, 10mM Tris [pH 8.0], 1mM EDTA, and 1x protease inhibitors) per 5×10^8^ parasites and (b) Sonicate the chromatin until sufficiently sheared (130µL, 5% duty cycle, 75W peak incident power, 200 cycles per burst, 7°C, for 5 minutes using Covaris Focus-Ultrasonicator M220). The immunoprecipitation step included: (a) Dilute the sample 1:5 with Dilution Buffer (0.01%v/v SDS, 1.1% TritonX-100, 1.2mM EDTA, 16.7mM Tris-HCl [pH 8.1], and 150mM NaCl), (b) Reduce background signal by pre-clearing with 20µL of Protein A/G magnetic beads (Millipore 16-663) per 1mL of sample for 2 hours at 4°C with rotation, (c) Aliquot 1/10 of the sample for the Input control and rotate at 4°C until DNA elution step, (d) Remaining 9/10 of sample is immunoprecipitated with 1:1000 anti-HA antibody (0.1mg/mL Roche Rat Anti-HA High Affinity [11867423001]) overnight at 4°C with rotation, (e) Collect the immune complexes with 20µL of Protein A/G magnetic beads per 1mL of sample for 2 hours at 4°C with rotation, (f) Wash (5 minutes each) immune complexes on magnet once with Low Salt Immune Complex Wash Buffer (0.1%v/v SDS, 1%v/v TritonX-100, 2mM EDTA, 20mM Tris-HCl [pH 8.1], and 150mM NaCl) at 4°C with rotation, once with High Salt Immune Complex Wash Buffer (0.1%v/v SDS, 1%v/v TritonX-100, 2mM EDTA, 20mM Tris-HCl [pH 8.1], and 500mM NaCl) at 4°C with rotation, once with LiCl Immune Complex Wash Buffer (0.25M LiCl, 1%v/v NP-40, 1%w/v Deoxycholate, 1mM EDTA, and 10mM Tris-HCl [pH 8.1]) at 4°C with rotation, and twice with 1xTE at room temperature with rotation, and (g) Elution from beads with 200µL fresh Elution Buffer (0.01%v/v SDS and 100mM NaHCO_3_) per 1mL of sample for 15 minutes at room temperature with rotation (also added to Input sample). The DNA purification step included: (a) Reverse-crosslink Input and IP samples with 0.2M NaCl and incubate overnight at 45°C with shaking (750rpm), (b) 0.4%v/v RNaseA treat from 30 minutes at 37°C with shaking (750rpm), (c) 0.3mg/mL Proteinase K treat for 2 hours at 45°C with shaking (750rpm), and (d) Purify DNA using MinElute column (Qiagen) as directed by manufacturer. Purified DNA samples were quantified using the Qubit fluorometer for high-sensitivity DNA.

### AP2-LT ChIP-seq library prep for Illumina sequencing

DNA sequencing libraries were prepared for high-throughput Illumina sequencing on the NextSeq 2000 with 150 x 150 single-end or paired-end mode. The library prep protocol includes 6 major steps: (1) end repair of DNA fragments, (2) addition of A-tail for adaptor ligation, (3) ligation of Illumina adaptors, (4) DNA sequence size selection, (5) DNA sequence amplification by PCR, and (6) DNA sequence clean up. The single indexed adaptors (Bioo Scientific) used were diluted 1:10 prior to ligation. DNA libraries were size selected for 250bp sequences. Due to the A/T-richness of the *P. falciparum* genome, KAPA HiFi polymerase was used for amplification during PCR steps. Only 12 rounds of amplification was used for AP2-LT ChIP-seq samples (both for input and IP libraries), and 16 rounds for the no-epitope negative control (both for input and IP libraries). After DNA sequence clean up, completed libraries were quantified using the Qubit fluorometer for high-sensitivity DNA and library sequence length by the Agilent Bioanalyzer 1000 or Agilent TapeStation 4150 before submitting for high-throughput sequencing.

### AP2-LT ChIP-seq data analysis and peak calling

Raw sequencing reads were first processed by trimming (Trimmomatic v0.32.3) Illumina adaptors and low quality reads below 30 Phred (SLIDINGWINDOW: 4:30). FastQC (v0.11.9) was used to check the quality after trimming. Processed reads were then mapped to the to the *P. falciparum* genome (release 38^79^) using BWA-MEM (v0.7.17.2) simple Illumina mode with multiple mapped reads filtered out (MAPQ=1). Once the sequences were mapped, MACS2^84^ was used to call peaks with each biological replicate and its paired input sample using a standard significance cutoff (q-value=0.01) (**Supplemental File 4**). Using BedTools Multiple Intersect (v2.29.2), the narrow peaks output file for each biological replicate was overlapped to identify the significant peaks in at least two of the three replicates (**Supplemental Figure 13C**). The overlapping regions were then used to identify an enriched DNA motif^85^ and target genes were defined by having a peak no more than 2kb upstream of the gene transcription start site (TSS)^86^ along with gene body peaks.

### Integrating gcPBM sequences with ChIP-seq analysis

The gcPBM sequences were mapped as detailed above with the ChIP-seq analysis. BedTools Multiple Intersect (v2.29.2) was used to identify motif-containing gcPBM sequences that were bound via ChIP-seq (“ChIP-bound”) by setting the minimum overlap at 58.3% of the 36-bp gcPBM sequences, to ensure the overlap contained the central motif (CACACA, GTGCAC, GTAC, TGCATGCA).

### Chromatin accessibility data analysis and comparison to gcPBM and ChIP-seq datasets

All sequences bound below the non-specific binding threshold were excluded and all gcPBM- bound sequences were further characterized into ChIP-bound vs. ChIP-unbound by overlapping the mapped gcPBM-bound sequences with ChIP-seq called peaks by MACS2. Overlap was determined using 400bp windows around the ChIP-seq peak midpoint and the gcPBM midpoint by using bedtools slop and bedtools intersect (bedtools version 2.30.0). Assay for Transposase-Accessible Chromatin using sequencing (ATAC-seq) data from Toenhake *et al*. 2018^44^ was used to assess chromatin accessibility. FASTQ files were downloaded with fastq-dump (version 2.9.6) from GEO (GSE104075) and then aligned to the *P. falciparum* 3D7 reference genome with bwa mem with -M (version 0.7.17). Aligned reads were filtered by mapping quality for >= 30 with samtools view (version 1.3.1). Then duplicates were removed with picard with commands MarkDuplicates-REMOVE_DUPLICATES true-VALIDATION_STRINGENCY STRICT (version 2.24.1). Each of the timepoints from Toenhake *et al.* 2018 had two ATAC-seq replicates. Additionally, Toenhake, *et al.* provided a control experiment (two replicates) consisting of Tn5-treated genomic DNA (gDNA). These controls were to account for the sequence bias of Tn5. The replicates were merged and then bedgraphs were created from the merged bams with bedtools genomecov with -bg flags and piped into sort-k1,1-k2,2n. These were smoothed with bedops--chop 100--stagger 100 and keeping the mean signal in the 100bp window with bedmap--mean (bedops and bedmap version 2.4.39). Each bin was normalized by dividing the bin’s value (representing the average signal in a 100bp window) by the number of mapped reads in the merged bam file. At the ChIP-seq peak midpoints, the ATAC-seq 100bp average was divided by the gDNA 100bp average signal after adding a 0.1 pseudocount to each bin to get the ratio of signal to control.

## Results

### Defining DNA-binding proteins that recognize similar DNA motifs or have overlapping genome-wide occupancies

To investigate the binding specificity of transcription factors (TFs) that recognize highly similar DNA sequence motifs, we selected a set of *P. falciparum* TFs with conserved DNA-binding domains (DBDs) and known position weight matrices (PWMs) representing related *in vitro* binding specificities (**Figure 1A**). The *P. falciparum* genome encodes four ApiAP2 proteins with putative paralogous AP2 domains which recognize a CACACA DNA motif^43^: PF3D7_0420300, PF3D7_0802100 (PfAP2-LT)^87, 88^, PF3D7_1456000 (PfAP2-HC)^71, 89^, and PF3D7_1305200 (**Figure 1A**). PfAP2-HC (Heterochromatin) has been extensively characterized and named for its localization to repressive heterochromatic regions via protein-protein interactions with heterochromatin protein 1 (PfHP1)^71, 89^. Additionally, PfAP2-LT (late trophozoite-expressed) was found to interact with PfGCN5 and PfPHD1, which are components of the PfSAGA (Spt–Ada–Gcn5 Acetyltransferase) transcriptional co-activator complex^87, 88^. PF3D7_0420300 and PF3D7_1305200 are thus far uncharacterized.

The *P. falciparum* genome also encodes three non-paralogous proteins that bind a GTGCAC DNA motif, which include two ApiAP2 proteins, PF3D7_0604100 (PfSIP2)^91, 92^ and PF3D7_1007700 (PfAP2-I)^83, 93^, as well as a homeodomain-like protein 1 (PF3D7_1466200; PfHDP1)^48^ (**Figure 1A**). PfSIP2 (SPE2-Interacting Protein) binds heterochromatic, sub-telomeric regions containing a bipartite SPE2-element DNA motif^91, 92^. PfAP2-I (Invasion), was identified to bind euchromatic, intergenic regions and was characterized to be a regulator of red blood cell invasion genes^83, 93^. Finally, PfHDP1 (Homeodomain-like Protein 1) plays a role in the early development of sexual stage parasites, called gametocytes, and is one of the first non-ApiAP2 sequence-specific TFs characterized in *P. falciparum*^48^.

We also investigated two additional TFs, PF3D7_1222600 (PfAP2-G)^83, 94^ and PF3D7_1466400 (PfAP2-EXP)^95^ (**Figure 1A**). PfAP2-G and PfAP2-EXP were included in this study to dissect the mechanisms driving the overlap of *in vivo* binding sites through TFs that recognize divergent *in vitro* DNA motifs. Based on comparisons of published *in vivo* binding site data from chromatin immunoprecipitation followed by sequencing (ChIP-seq) studies, PfAP2-G and PfAP2-EXP recognize unique DNA motifs yet occupy a subset of overlapping genomic regions with PfAP2-I^83, 95^. PfAP2-G (Gametocytogenesis) is a master regulator of sexual stage commitment and binds a GTAC DNA motif both *in vitro* and *in vivo*^43, 83, 94^*. Pf*AP2-EXP (Exported Proteins) recognizes a TGCATGCA DNA motif through *in vitro* and *in vivo* binding assays, including a high-resolution structural study^62, 95, 96^. While overlapping genomic binding sites have been reported for a number of TFs in *P. falciparum*^71, 83, 95, 97^, the mechanisms that drive these shared binding events are not well understood.

For ApiAP2 TFs with multiple AP2 DBDs (PF3D7_0420300 [two domains], PfSIP2 [two domains], and PfAP2-I [three domains]) we selected only the AP2 domain which recognizes the motif-of-interest (**Figure 1A**; **Supplemental Figure 1A**). Amino acid sequence alignments for three of the four CACACA-binding AP2 DBDs showed a high percent identity (33.33-49.02%), with a lower percent identity (21.88%) among the GTGCAC-binding AP2 DBDs (**Supplemental Figure 1B,C**). The CACACA-binding DBD from PF3D7_1305200 was less conserved relative to the other CACACA-binding DBDs (20.37% identity) (**Supplemental Figure 1B**) and is not expressed during the asexual blood stage^98^. Therefore, PF3D7_1305200 was excluded from this study. Overall, the binding specificities for eight DBDs were investigated herein.

### High throughput interrogation of all possible transcription factor binding sites by gcPBM

To determine the full spectrum of *in vitro* binding specificities across the *P. falciparum* genome for the eight candidate DBDs, we designed and synthesized a novel *P. falciparum*-specific genomic-context protein-binding microarray (gcPBM) (**Figure 1B**). This gcPBM design allowed for simultaneous examination of all possible genome-wide motif occurrences for each of the eight DBDs flanked by *P. falciparum* genomic sequence context. Using PWM data from previously published work^42, 43^, we identified all instances of the DNA motifs, centered on a 36-bp window across all intergenic regions of the *P. falciparum* genome (PlasmoDB^79^: 3D7 strain genome release v38; motif E-score cutoff of > 0.45). In addition, each motif type (CACACA, GTGCAC, GTAC, and TGCATGCA) bound by the eight DBDs had associated non-motif-containing negative control DNA probes represented on the gcPBM (**Figure 1B**). The negative control probes consisted of randomly selected unique genomic sites lacking the motif-of-interest to assess non-specific binding of each DBD to the A/T-rich *P. falciparum* genome. After discarding redundant sequences, this resulted in: 9388 CACACA probes (1834 negative controls), 1394 GTGCAC probes (736 negative controls), 8998 GTAC probes (620 negative controls), and 1059 TGCATGCA probes (612 negative controls), which together represented 24,641 unique 36-bp genomic sequences (**Figure 1B; Supplemental File 1**).

Each unique DNA sequence was represented in both a 5′ or 3′ orientation, with one end of each DNA molecule attached to the glass slide (**Supplemental Figure 2A**). Additionally, each DNA probe was replicated randomly on the gcPBM (four CACACA/GTGCAC replicates and three GTAC/TGCATGCA replicates per orientation). Replicate probes were used to calculate the median signal intensity from both probe orientations. Overall, the total number of dsDNA probes was 174,550 spots, which were arrayed using an Agilent Technologies 4×180k microarray design. This *P. falciparum*-centric gcPBM was designed to identify the protein-DNA specificity in a high-throughput manner for each DBD across all intergenic instances of their cognate motif, allowing for a comprehensive interrogation of the impact of *P. falciparum* genomic context on TF binding.

### CACACA-binding transcription factors modestly prefer differing DNA sequence context in vitro

Candidate DBDs were tested on the gcPBM in addition to a technical replicate of a representative DBD per motif type (*i.e.,* AP2-LT for the CACACA group). After calculating the median binding intensity for the replicates of each orientation, we observed differences between the binding intensity values (**Supplemental Figure 2B-M**). Therefore, we selected the highest natural log median binding intensity value from either the 5′ or 3′ orientation for each unique DNA probe. Given the degree of amino acid identity (33.33-49.02%) between the three CACACA- binding AP2 domains (**Supplemental Figure 1B**), we used the gcPBM to directly test if there were divergent sequence preferences for binding to the CACACA motif when surrounded by differing genomic contexts. As expected, each binding experiment demonstrated a significant preference for CACACA probes over associated negative control probes (Two-tailed Mann-Whitney test [p-value < 0.0001]) (**Figure 2A**; **Supplemental Figure 3A-C**). However, the distribution of binding to CACACA probes was narrow as compared to the negative control distribution, indicating that sequence context had limited importance to binding specificity (**Figure 2A**; **Supplemental Figures 3A-C**). For each CACACA-binding DBD, the top 100 probes bound at the highest gcPBM binding intensities showed a strong preference towards long CA- dinucleotide repeats, with a slightly degenerate AT-dinucleotide repeat in the 3′ flanks (**Figure 2B**, **Supplemental Figure 3A-C**). Individual pairwise comparisons of the binding intensities between all three of the paralogous CACACA-binding DBDs showed only moderate differences in DNA sequence preference: R^2^ = 0.80 (PF3D7_0420300_D1 vs. AP2-LT), R^2^ = 0.86 (PF3D7_0420300_D1 vs. AP2-HC), and R^2^ = 0.85 (AP2-LT vs. AP2-HC), respectively (**Figure 2D-F**) versus the AP2-LT technical replicate experiment (R^2^ = 0.95) (**Figure 2C**). These similarities in high affinity sites, along with the modest differences in sequence context preferences, suggested to us that DNA binding preferences diverge at lower affinity binding.

**Figure 2:**
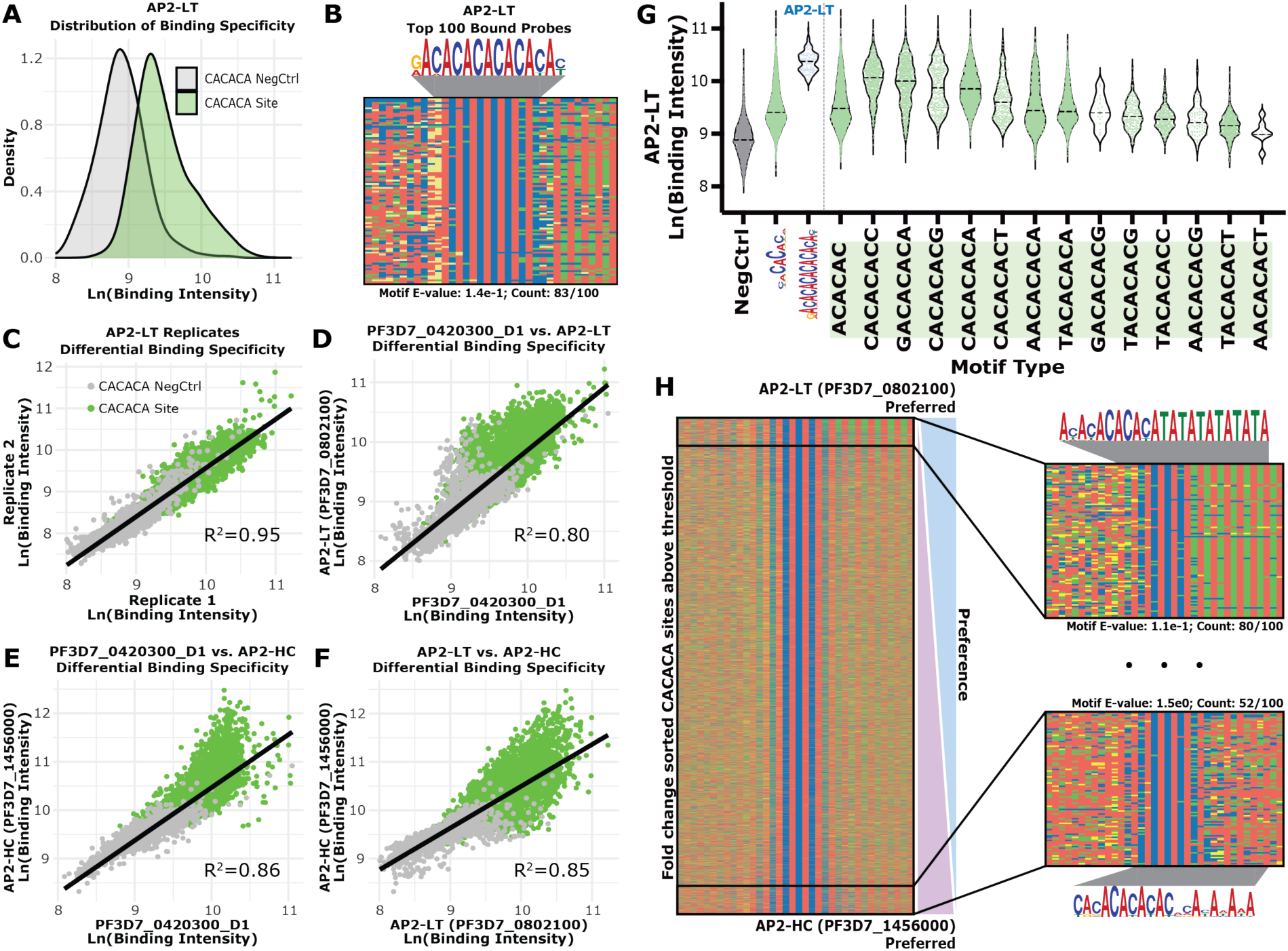
CACACA-binding AP2 domains have moderate differences in sequence context preferences at medium-to-low affinities **(A)** Binding intensity distributions for CACACA probes and respective negative control probes for AP2-LT (CACACA probes [Green] and negative control probes [Grey]); **(B)** Four-color plot of the top 100 bound probes by AP2-LT with enriched motif above and calculated E-value and number of occurrences below^85^. Color representations: A (Red), C (Blue), G (Yellow), and T (Green); **(C)** Comparison of the gcPBM binding intensities for AP2-LT technical replicates (Pearson correlation: R^2^=0.95) (CACACA probes [Green] and negative control probes [Grey]); **(D)** Comparison of the gcPBM results for PF3D7_0420300_D1 versus AP2-LT (Pearson: R^2^=0.80); **(E)** Comparison of the gcPBM results for PF3D7_0420300_D1 versus AP2-HC (Pearson: R^2^=0.86); **(F)** Comparison of the gcPBM results for AP2-LT versus AP2-HC (Pearson: R^2^=0.85); **(G)** Binding intensity distributions from AP2-LT for negative control probes (Grey), all CACACA probes (Green), the AP2-LT extended motif probes (Blue), and all 8-mer CACACA probes represented in the gcPBM (Green; right of the vertical line). Dotted lines in each violin plot are the calculated mean; **(H)** *Left:* Four-color plot of CACACA probes above the 90^th^ percentile of negative control probes sorted by fold change (log2[AP2-LT/AP2-HC]). *Right:* Zoom in on the top 100 differentially bound sites by AP2-LT (*top right*) and AP2-HC (*bottom right*) with enriched motifs, calculated E-values, and motif occurrence counts within those top 100 sites^85^.

To determine whether the high prevalence of longer CA-dinucleotide repeats was due to an actual binding preference or because the long CA-repeats created multiple binding events per DNA probe, we used electrophoretic mobility shift assays (EMSAs). Systematic mutation of two of the seven CA-dinucleotide repeats to TT-dinucleotides across a representative high affinity CACACA probe abolished the slower/higher-mobility shift (multimeric binding of AP2-LT DBDs) upon mutation of two central CA-dinucleotide repeats (**Supplemental Figure 4A**). This suggests that many of the AP2-LT high affinity probes with more than three CA-dinucleotide repeats likely resulted from an interaction between more than one AP2-LT DBD per DNA probe. EMSAs with the same DNA probes against AP2-HC DBD did not result in a slower-mobility shift demonstrating that AP2-HC likely interacts with the longer repeats in a 1:1 (DBD:DNA) stoichiometry, unlike the AP2-LT DBD (**Supplemental Figure 4A**).

Due to the apparent multimeric binding to high affinity probes by AP2-LT, we expanded our analysis to include all CACACA probes (above negative control signal) to further explore preferences for nucleotides directly adjacent to the central CACACA motif at various levels of affinity. To categorize each protein-DNA binding event into specifically bound versus non-specifically bound, we set a threshold at the 90^th^ percentile of the binding signal for negative control probes, as per previous studies^10, 12^. To explore the distributions of binding intensities within one nucleotide on either side of the central 6-mer, we parsed out the binding intensities for each 8-mer sequence represented on the gcPBM with the “ACACAC” as the most represented central 6-mer (**Supplemental File 5**). We found that the central 8-mer sequences with the highest average binding signal were all extensions of the CA-dinucleotide repeat (**Figure 2G**; **Supplemental Figure 5**). However, calculating the fold change of the binding intensities between each pairwise comparisons (above the negative control threshold) showed a strong preference for a short CA-repeat with flanking AT-dinucleotide repeats for AP2-LT, extended CA-repeats (up to six CA-dinucleotide repeats) for AP2-HC, and short CA-repeats with no flanking pattern for PF3D7_0420300_D1 (**Figure 2H**; **Supplemental Figure 6**). These differences in sequence context at lower affinity probes suggest that, although the CACACA-binding DBDs did not prefer strikingly different sequence contexts at the high affinity probes, the modest differences in binding resulted from the medium-to-low affinity ranges.

Additional EMSA-based validation using AP2-LT and gcPBM dsDNA probes with low-, medium-, and high-affinities from both CACACA and negative control probes recapitulated the varying degrees of binding specificities shown in the high-throughput gcPBM experiments (**Supplemental Figure 4B)**. Overall, these findings demonstrate that DNA-binding of the CACACA-binding ApiAP2 DBDs is not greatly impacted by sequence context and these proteins differ in their abilities to multimerize on DNA *in vitro*.

### GTGCAC-binding transcription factors have distinct sequence context preferences by gcPBM

Based on the available *in vivo* ChIP-seq genome-wide binding sites for the GTGCAC- binding TFs, PfAP2-I^83^ and PfHDP1^48^, we found many uniquely bound sites with only 47 shared sites (**Supplemental Figure 7A**). Therefore, to investigate whether DNA sequence context explains differential binding site selection, we tested GTGCAC-binding DBDs using our *P. falciparum* gcPBM. The GTGCAC-binding group includes one AP2 domain each from two different ApiAP2 proteins (domain one [D1] from SIP2 and domain three [D3] from AP2-I) and a homeodomain from HDP1 (**Figure 1A**). All three GTGCAC-binding DBDs tested preferred GTGCAC probes over negative control probes (Two-tailed Mann-Whitney test [p-value < 0.0001]) (**Figure 3A**; **Supplemental Figure 3D-F**). In contrast to the paralogous CACACA- binding DBDs (**Figure 2A**), the distribution of binding to GTGCAC probes was much broader than the respective negative control distribution, which suggested a greater impact of sequence context on DNA binding (**Figure 3A**; **Supplemental Figure 3D-F**). Additionally, the top 100 GTGCAC probes bound by the three DBDs revealed a more extended sequence preference beyond the previously published 6-mer DNA motifs^43, 48^, such as GGTGCAC for SIP2_D1, AGTGCATTA for AP2-I_D3, and TGTGCACA for HDP1 (**Figure 3B**; **Supplemental Figure 3D-F**).

**Figure 3:**
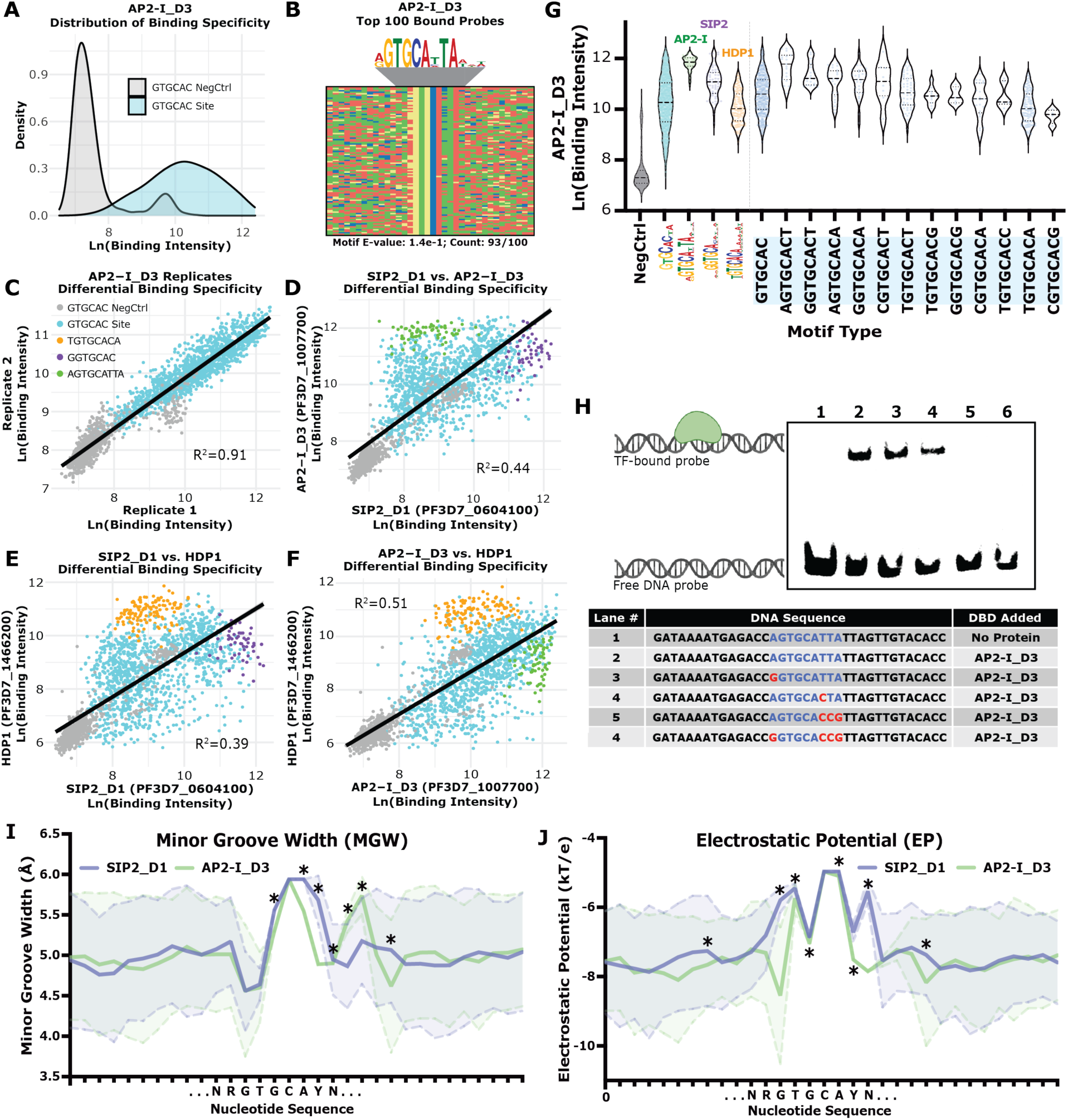
Binding specificity is dependent on nucleotides proximal to the GTGCAC motif **(A)** Binding intensity distributions for GTGCAC probes and the respective negative control probes for AP2-I_D3. (GTGCAC probes [Blue] and negative control probes [Grey]); **(B)** Four-color plot of top 100 bound probes by AP2-I_D3 with enriched motif above and calculated E-value and number of occurrences below^85^. Color representations: A (Red), C (Blue), G (Yellow), and T (Green); **(C)** Comparison of the binding intensities for AP2-I_D3 technical replicates (Pearson correlation: R^2^=0.912). (Negative control probes [Grey], GTGCAC probes [Blue], HDP1- preferred TGTGCACA probes [Orange], SIP2_D1-preferred GGTGCAC probes [Purple], and AP2-I_D3-preferred AGTGCATTA probes [Green]; **(D)** Comparison between SIP2_D1 and AP2-I_D3 (Pearson: R^2^=0.442). **(E)** Comparison between SIP2_D1 and HDP1 (Pearson: R^2^=0.386). **(F)** Comparison between AP2-I_D3 and HDP1 (Pearson: R^2^=0.514); **(G**) Binding intensity distributions from AP2-I_D3 for GTGCAC negative control probes (Grey), all GTGCAC probes (Blue), the extended motif probes by all three GTGCAC-binding TFs (AP2- I_D3[Green], SIP2_D1[Purple], and HDP1[Orange]), and 8-mer GTGCAC probes represented in the gcPBM (Blue; right of the vertical line); **(H)** EMSA of AP2-I_D3 binding to a AGTGCATTA probe with increasing numbers of mutations to the extended motif. Protein-DNA interaction graphic generated using BioRender; **(I)** Calculated minor groove width (MGW) predictions across all AGTGCATTA probes (Green) and all GGTGCAC probes (Purple)^82^. * Denotes statistically significant differences (p-value < 0.05) between means (two-sided Wilcoxon rank sum test). N = IUPAC for any nucleotide. Y = IUPAC for C or T nucleotides; **(J)** Calculated electrostatic potential (EP) predictions across all AGTGCATTA probes (Green) and all GGTGCAC probes (Purple)^82^. * Denotes statistically significant differences (p-value < 0.05) between means (two-sided Wilcoxon rank sum test). N = IUPAC for any nucleotide. Y = IUPAC for C or T nucleotides.

Individual pairwise comparisons of the gcPBM data between all three GTGCAC DBDs demonstrate pronounced differences in DNA sequence context preferences (**Figure 3D-F**) with low Pearson correlations: R^2^ = 0.44 (SIP2_D1 vs. AP2-I_D3), R^2^ = 0.39 (SIP2_D1 vs. HDP1), and R^2^ = 0.51 (AP2-I_D3 vs. HDP1), respectively (**Figure 3D-F**), compared to that of the AP2- I_D3 technical replicates (R^2^ = 0.91; **Figure 3C**). To broadly identify sequence preferences proximal to the central 6-mer GTGCAC DNA motif, the binding intensity for each GTGCAC- binding DBD was compared across the three extended motifs (AP2-I_D3: AGTGCATTA, SIP2_D1: GGTGCAC, and HDP1: TGTGCACA) (**Figure 3G**; **Supplemental Figure 8**). As expected, each DBD showed a significant preference towards binding the extended motifs we identified. We found additional preferences for other single nucleotides proximal to the GTGCAC core sequence by parsing out the binding intensities of all 8-mers with central “GTGCAC” sequences represented on the gcPBM (**Supplemental File 5**), which further indicated an influence of sequence context on TF binding *in vitro* for the GTGCAC-binding TFs (**Figure 3G; Supplemental Figure 8**). While there are clear differences in the top 100 probes bound by each GTGCAC-binding DBD (**Figure 3B**, **Supplementary Figures 3D-F**), by calculating the fold change between binding intensities for each pairwise comparison, we failed to identify any secondary extended motifs compared to those identified above (**Supplemental Figure 9**).

By EMSA, we tested whether the extended motif bound by AP2-I_D3 was necessary for DNA binding (**Figure 3H**). The dsDNA probe used was a representative 36-bp sequence which contained the central AGTGCATTA extended motif, was bound at a high affinity in the *in vitro* gcPBM, and was located in an *in vivo* AP2-I ChIP-seq bound region^83, 93^. Increased mutations to the extended nucleotides flanking the core GTGCAC 6-mer motif caused a reduction in AP2-I_D3 binding, demonstrating that the extended DNA sequence preferred by AP2-I_D3 is important for binding (**Figure 3H**). Overall, these results indicate that the sequence context of the GTGCAC motif greatly influences differential binding between the SIP2_D1, AP2-I_D3, and HDP1 GTGCAC-binding DBDs.

### GTGCAC-binding transcription factors prefer DNA sequences with diverse predicted DNA shapes

We noted that specific nucleotide patterns distal to the GTGCAC motif (more than three nucleotides upstream/downstream of the extended motif) were not enriched in the probes preferred by the GTGCAC-binding factors (**Supplemental Figure 9**). Therefore, we explored whether “shape/indirect-readout” of sequence-dependent DNA topologies might contribute to binding in addition to base-specific DBD contacts through “base/direct-readout” mechanisms^5^. Predicted DNA shape measurements take into consideration how the local flexibility or intrinsic shape of the DNA impacts docking of a TF into its preferred DNA motif and includes the effects of 2- to 5- mer nucleotide patterns via inter- and intra-nucleotide interactions on TF binding specificity. To identify the possible impact of flanking nucleotide patterns beyond the extended GTGCAC DNA motif, we predicted intrinsic DNA shape features for the extended motif-containing probes. While there are numerous DNA shape features^5^, we used minor groove width (MGW) and electrostatic potential (EP), which have been shown previously to be highly predictive of DNA-binding specificity^81, 99–101^. Since the CACACA group did not prefer divergent extended motifs (**Figure 2B**, **Supplemental Figure 3A-C**), we focused on DNA shape analysis on the GTGCAC group. Using DNAShapeR^82^, we predicted MGW and EP profiles for all DNA probes containing the AGTGCATTA, GGTGCAC, and TGTGCACA extended motifs. Significant differences (two-sided Wilcoxon rank sum test; p-value < 0.05) between each pairwise comparison for MGW and EP suggested that, in addition to base-specific contacts, DNA shape-readout mechanisms may also influence *in vitro* binding specificity of the GTGCAC-binding DBDs (**Figure 3I,J; Supplemental Figure 10A-D**).

To further probe the potential impact of our predicted DNA shape on binding, the AP2- I_D3:DNA interaction was tested by EMSA using specific DNA mutations that maximize the change of predicted MGW and EP (*i.e.,* shape-readout mutations), while minimizing the number of mutated nucleotides (**Supplemental File 3**). Distal shape mutations (more than three nucleotides upstream/downstream of the extended AGTGCATTA motif) (**Supplemental Figure 11**) did not impact the strength of the AP2-I_D3:DNA bound state (**Supplemental Figure 10E**), suggesting that specific DNA shapes of the distal flanking sequences are not required for AP2- I_D3 to bind a high-affinity site. Therefore, we conclude that DNA sequence/shape context of proximal, but not distal, nucleotides relative to the GTGCAC motif may contribute to differential specificity for the GTGCAC-binding DBDs.

### CACACA-binding transcription factors occupy differential sites in vivo

Since the binding specificity of the CACACA-binding DBDs were not greatly impacted by sequence context, we investigated the potential impact of the *in vivo* chromatin landscape on genome-wide binding site selection. While AP2-HC has been fully characterized *in vivo*^89^, the other CACACA-binding TFs, PF3D7_0420300 and AP2-LT, have not. Therefore, we determined the *in vivo* genome-wide occupancy of AP2-LT using ChIP-seq from *P. falciparum* cell culture. The endogenous locus of *pfap2-lt* was modified using CRISPR-Cas9 to insert a sequence encoding a C-terminal (3x)hemaglutininA (HA) tag (**Supplemental Figure 12**) to generate AP2-LT^HA^. Three biological replicates of ChIP-seq using antibodies against the HA tag from a clonal parasite line of AP2-LT^HA^ and a negative control no-epitope sample were conducted at the peak of AP2- LT protein expression during the early schizont stage (36-45 hours post invasion [hpi]) (**Supplemental Figure 13**)^98^. While our *in vitro* binding assays suggested that AP2-LT preferred a CACACA motif, our ChIP-seq results surprisingly identified that the most enriched bound DNA sequence motif^85^ was TGCAC (E-value=4.6e-190) (**Figure 4A**). In addition, we also found the occurrence of a longer motif TGCACN_5_TGCAC (E-value=9.0e-272) (**Figure 4B**; **Supplemental Figure 14**) that contained two TGCAC motifs starting at position one and position eleven. This bipartite motif therefore encompasses a full turn of the DNA helix, suggesting potential AP2-LT dimerization as we found via EMSA (**Supplemental Figure 4B**). The AP2-LT *in vivo*-bound TGCAC motif is quite different than the *in vitro*-bound CACACA motif and surprisingly resembles the GTGCAC motif bound by SIP2_D1, AP2-I_D3, and HDP1. We hypothesize that specific features of the nuclear environment such co-factor interactions, multimerization, chromatin accessibility, and/or epigenetic marks may contribute to the divergent *in vivo* sequence specificity for AP2-LT.

**Figure 4:**
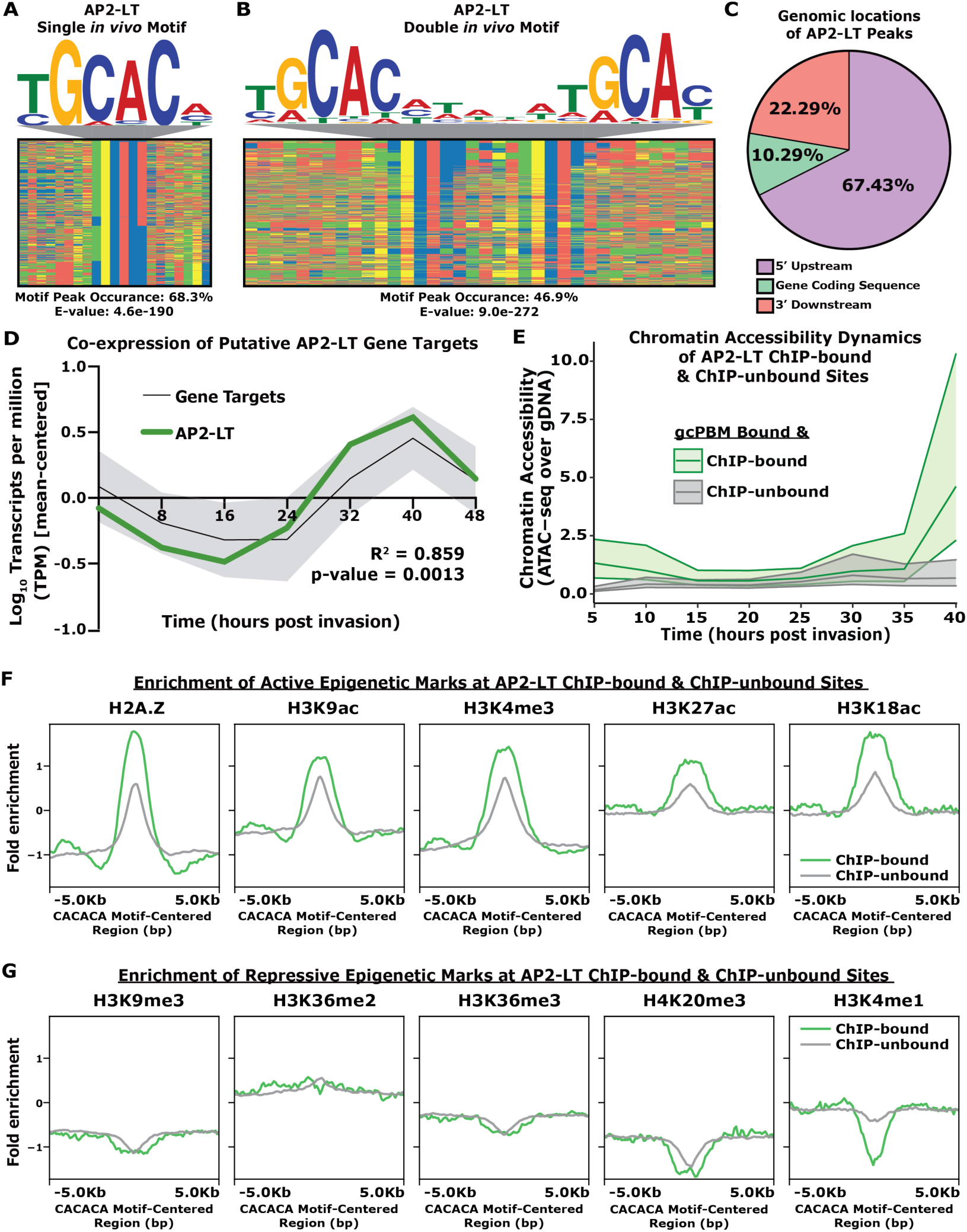
AP2-LT mostly binds to intergenic regions upstream of late-stage genes demarcated by chromatin accessibility and active epigenetic modifications *in vivo* **(A)** Four-color plot of TGCAC-centered AP2-LT bound sites, enriched DNA motif above, calculated motif peak occurrence, and calculated motif E-value below^85^. Color representations: A (Red), C (Blue), G (Yellow), and T (Green); **(B)** Four-color plot of TGCACN5TGCAC-centered AP2-LT bound sites, enriched DNA motif above, calculated motif peak occurrence, and calculated motif E-value below^85^; **(C)** Percent of MACS2-called peaks that overlap with 5‵- upstream regions (Purple), gene coding sequences (Green), or 3‵- downstream regions (Peach); **(D)** Mean-centered transcript abundance profile of 275 AP2-LT gene targets (mean [Black] with one standard deviation [Grey]) compared to the mean-centered transcript abundance profile of the AP2-LT transcript (Bold Green). Calculated R^2^ and p-value from Pearson correlation between putative gene targets and AP2-LT profiles bottom right; (**E**) Chromatin accessibility across eight asexual stage timepoints (5hpi, 10hpi, 15hpi, 20hpi, 25hpi, 30hpi, 35hpi, and 40hpi) for AP2-LT ChIP-bound (Green) and ChIP-unbound (Grey) sites. Central line plotted is the median normalized read count over gDNA control. Upper and lower lines are the 75^th^ percentile and 25^th^ percentile, respectively; (**F**) Profile plot of the mean ChIP-seq fold enrichment (Log2[IP/Input]) of five active epigenetic marks (H2A.Z, H3K9ac, H3K4me3, H3K27ac, and H3K18ac) for ChIP-bound (Green) and ChIP-unbound (Grey) sites; and (**G**) Profile plot of the mean ChIP-seq fold enrichment (Log2[IP/Input]) of five repressive epigenetic marks (H3K9me3, H3K36me2/3, H4K20me3, and H3K4me1) for ChIP-bound (Green) and ChIP-unbound (Grey) sites.

Our ChIP-seq data indicated that AP2-LT mostly occupies regions upstream of transcription start sites (TSSs) as well as gene coding sequences (**Figure 4C; Supplemental File 4**). Putative target genes (**Supplemental File 6**) were predicted based on the presence of a ChIP-seq peak no more than 2kb upstream of a target gene TSS^86^ or within the gene coding sequence. When a peak was located within 2kb upstream of head-to-head genes, the closest gene was selected. In general, the transcript abundance profiles for these putative target genes revealed that they are co-expressed with the AP2-LT transcript (**Figure 4D**), with GO terms^79^ associated with parasite egress/invasion, protein modifications, and cell cycle (**Supplemental File 7**). These findings are consistent with AP2-LT acting as an activator of late-stage genes, which is further supported by previous reports that AP2-LT is a component of the PfSAGA co-activator complex^87, 103^.

We find that AP2-LT and AP2-HC bind mutually exclusive genome-wide binding sites *in vivo* (**Supplemental Figure 7B**). Our data show that AP2-LT primarily binds in euchromatic regions, while previous work found AP2-HC mostly localized to heterochromatin via interactions with HP1^89^. Overall, AP2-LT and AP2-HC do not occupy the same sites *in vivo* likely due to the influence of the nuclear environment, including the capacity to multimerize and co-factor interactions.

### Chromatin accessibility and histone modifications differentiate in vivo binding site selection of CACACA-binding transcription factors

To further probe whether CACACA sequence context impacts TF binding site selection in the context of the nuclear environment, we compared genomic regions bound by gcPBM versus those bound in the ChIP-seq data. The gcPBM data was first grouped into gcPBM-bound and gcPBM-unbound probes using a threshold set at the 90^th^ percentile of the CACACA negative control probes. The gcPBM-unbound probes (“non-specific binding”) were excluded. The gcPBM-bound probes were further grouped into ChIP-bound vs. ChIP-unbound sites using a minimum overlap of 21-bp of the 36-bp gcPBM sequences to ensure that the overlap contained the central motif of the mapped gcPBM-bound sequences within ChIP-seq called peaks by MACS2 (**Supplemental File 4**). Interestingly, AP2-LT ChIP-bound sites were bound *in vitro* at significantly higher gcPBM binding intensities than all the CACACA sites, while the few AP2- HC ChIP-bound sites (207) were bound significantly lower than all the CACACA sites (**Supplemental Figure 15A,B**). These results demonstrate that high affinity *in vitro* DNA-binding preference alone does not predict binding site selection *in vivo* for the CACACA-binding TFs.

Predicting DNA motifs for the ChIP-bound and ChIP-unbound sites also showed that the longer CA-dinucleotide repeats and flanking AT-repeats found by gcPBM (**Figure 2H**) were not enriched in the AP2-LT ChIP-bound sites (**Supplemental Figure 15A**), further suggesting a shift in sequence preference between *in vitro* and *in vivo* binding for AP2-LT. After reanalyzing the published AP2-HC ChIP-seq^89^ data, several euchromatic peaks contained the CACACA motif although it was not enriched overall (**Supplemental Figure 15B**). Additionally, we found a change from a “CACACA” preference *in vitro* to “TGCAC” *in vivo* by AP2-LT (**Supplemental Figure 16A,B**) and a low Pearson correlation between *in vitro* and *in vivo* binding for AP2-LT (**Supplemental Figure 16C**) and AP2-HC (**Supplemental Figure 16D**). These findings suggested that *in vitro-*defined sequence context preferences are not the main drivers of *in vivo* binding site selection for AP2-LT and AP2-HC.

To investigate the possible contribution of the chromatin landscape on TF binding, we compared the ChIP-bound and ChIP-unbound sites with published temporal genome-wide chromatin accessibility and epigenetic post-translational modifications (PTMs) datasets. For this analysis, we used the ChIP-unbound sites as a control to observe if changes in chromatin accessibility and epigenetic patterns were unique to the ChIP-bound sites. Using reanalyzed Assay for Transposase-Accessible Chromatin using sequencing (ATAC-seq) data^44^ from eight timepoints, we found that ChIP-unbound sites were largely inaccessible throughout the 48-hour asexual developmental cycle (**Figure 4E**; **Supplemental Figure 17)**. More interestingly, a dynamic opening of chromatin was observed at the ChIP-bound sites only during late-stage development, which coincides with the maximal transcript and protein expression of AP2-LT^98, 104^ (**Figure 4E**; **Supplemental Figure 17**). In contrast, AP2-HC associates with regions of inaccessible chromatin throughout parasite development, as expected (**Supplemental Figure 15B**).

We next determined whether ChIP-bound and ChIP-unbound sites were differentially demarcated by histone activation or repression marks^67–69^ in schizonts when AP2-LT and AP2-HC are maximally expressed^98, 104^. We found that activation marks (H2A.Z, H3K9ac, H3K4me3, H3K27ac, and H3K18ac) were highly enriched at AP2-LT ChIP-bound sites, while the repression marks (H3K9me3, H3K36me2, H3K36me3, H4K20me3, and H3K4me1) were not (**Figure 4F,G**; **Supplemental Figure 18)**. In contrast, AP2-HC ChIP-bound sites were enriched with repressive marks at binding sites, with the most represented mark being heterochromatic H3K9me3, as expected (**Supplemental Figure 19**). We also found only a modest difference of epigenetic mark enrichment between the ChIP-bound versus ChIP-unbound sites, which suggests that these epigenetic marks co-occur at ChIP-bound sites, but do not define the binding site selection between ChIP-bound and ChIP-unbound sites. While single cell RNA-seq (scRNA-seq) data suggests that AP2-LT and AP2-HC are co-expressed in almost all cells during the asexual blood stage^74^ (**Supplemental Figure 20A-D**), these results demonstrate that the mutually exclusive genomic target site selection by AP2-LT and AP2-HC are impacted by chromatin state.

### Sequence context, chromatin state, and timing of expression differentiate in vivo binding site selection of GTGCAC-binding transcription factors

As demonstrated by the gcPBM binding results (**Figure 3**), DNA sequence context and intrinsic DNA shape can impact the *in vitro* binding of the GTGCAC-binding DBDs (SIP2_D1, AP2-I_D3, and HDP1). To explore the impact of sequence context on binding site selection of the GTGCAC group *in vivo*, we compared the *in vitro* gcPBM binding intensities to the *in vivo* ChIP-seq genome-wide occupancy data for AP2-I^83^ and HDP1^48^ due to their public availability. As found by gcPBM (**Figure 3B**), genomic sites bound by AP2-I *in vivo* also revealed an enrichment for the extended AGTGCATTA motif (**Supplemental Figure 15C**; **Supplemental Figure 21A-C**). To identify the impact of chromatin context, we compared ATAC-seq chromatin accessibility^44^ and ChIP-seq epigenetic PTMs^67–69^ to AP2-I ChIP-bound and ChIP-unbound sites. Our results showed ChIP-bound sites were almost exclusively accessible, while the ChIP-unbound sites were largely inaccessible (**Supplemental Figure 15C**). Comparisons to epigenetic PTMs showed an enrichment of all five activation marks tested (H2A.Z, H3K9ac, H3K4me3, H3K27ac, and H3K18ac) (**Supplemental Figure 22**), suggesting that AP2-I likely requires chromatin-accessible sites containing activating epigenetic PTMs.

A reanalysis of HDP1 ChIP-seq data identified genome-wide binding to intergenic regions upstream of genes during sexual blood stage development^48^. We found that the extended TGTGCACA motif preferred by HDP1 *in vitro* (**Supplemental Figure 3F**) is also enriched *in vivo* (**Supplemental Figure 15D**), although there was low correlation between the *in vitro* and *in vivo* bound sites (**Supplemental Figure 21D-F**). While scRNA-seq data suggests AP2-I and HDP1 are only occasionally co-expressed in individual cells during the asexual blood stage^74^ (**Supplemental Figure 20E-H**), distinct *in vivo* binding site selection is largely due to differences in the extended DNA sequence motifs.

### Low affinity DNA-binding preferences across divergent DNA motifs may influence TF co-occupancy genome-wide

Because the *P. falciparum*-specific gcPBM design contains all four motif types (CACACA, GTGCAC, GTAC, and TGCATGCA) (**Figure 1B**) this allowed us to reanalyze our data to interrogate TF binding to other DNA motifs on the array. Previous work has shown that some ApiAP2 TFs that recognize different DNA motifs can occupy overlapping genomic regions^71, 83, 105^. Two such factors, AP2-G (GTAC-binder) (**Figure 1A**) and AP2-I (GTGCAC- binder), share roughly one-third of *in vivo* binding sites by ChIP-seq (**Supplemental Figure 7C**). Moreover, single-cell transcriptomics^74^ demonstrates that AP2-G and AP2-I (**Supplemental Figure 20I,J**) are co-expressed in a limited subset of cells.

To characterize possible mechanisms of cooperative binding between AP2-G and AP2-I we first determined the genome-wide binding specificity of AP2-G using the *P. falciparum*- specific gcPBM. AP2-G bound to GTAC probes with a narrow distribution of binding and high correlation between technical replicates (**Supplemental Figure 23A-D)** which implied a low importance for sequence context. While the AP2-G ChIP-bound GTAC sites resulted in higher binding intensities over all other GTAC sites (**Supplemental Figure 15E**), the *in vitro* gcPBM versus *in vivo* ChIP-seq binding were only moderately correlated (**Supplemental Figure 24A-C**). We then analyzed the AP2-G and AP2-I_D3 gcPBM data for binding to both GTAC- and GTGCAC-containing probes. Interestingly, AP2-G bound GTGCAC probes significantly above GTAC negative control probes (**Figure 5A**) and AP2-I_D3 bound GTAC probes significantly above GTGCAC negative control sites (**Figure 5B**) at low affinities, suggesting that these DBDs have the capacity to bind divergent DNA motifs *in vitro*. To determine if these low affinity sites are also bound *in vivo* by both AP2-G and AP2-I, we next compared the AP2-G and AP2-I_D3 ChIP-bound sites to categorize sites that were co-occupied by both TFs (“co-ChIP-bound”). We calculated the fold change of gcPBM binding between the co-ChIP-bound sites and found a portion that are preferred by AP2-I (Log_2_FC > 2), by AP2-G (Log_2_FC >-2), or were equally preferred by both DBDs *in vitro* (2 > Log_2_FC >-2) (**Figure 5C**). These results show that the AP2 domains of AP2-G and AP2-I_D3 can bind to co-occupied sites at low affinities, which may influence the co-occupancy observed *in vivo*.

**Figure 5:**
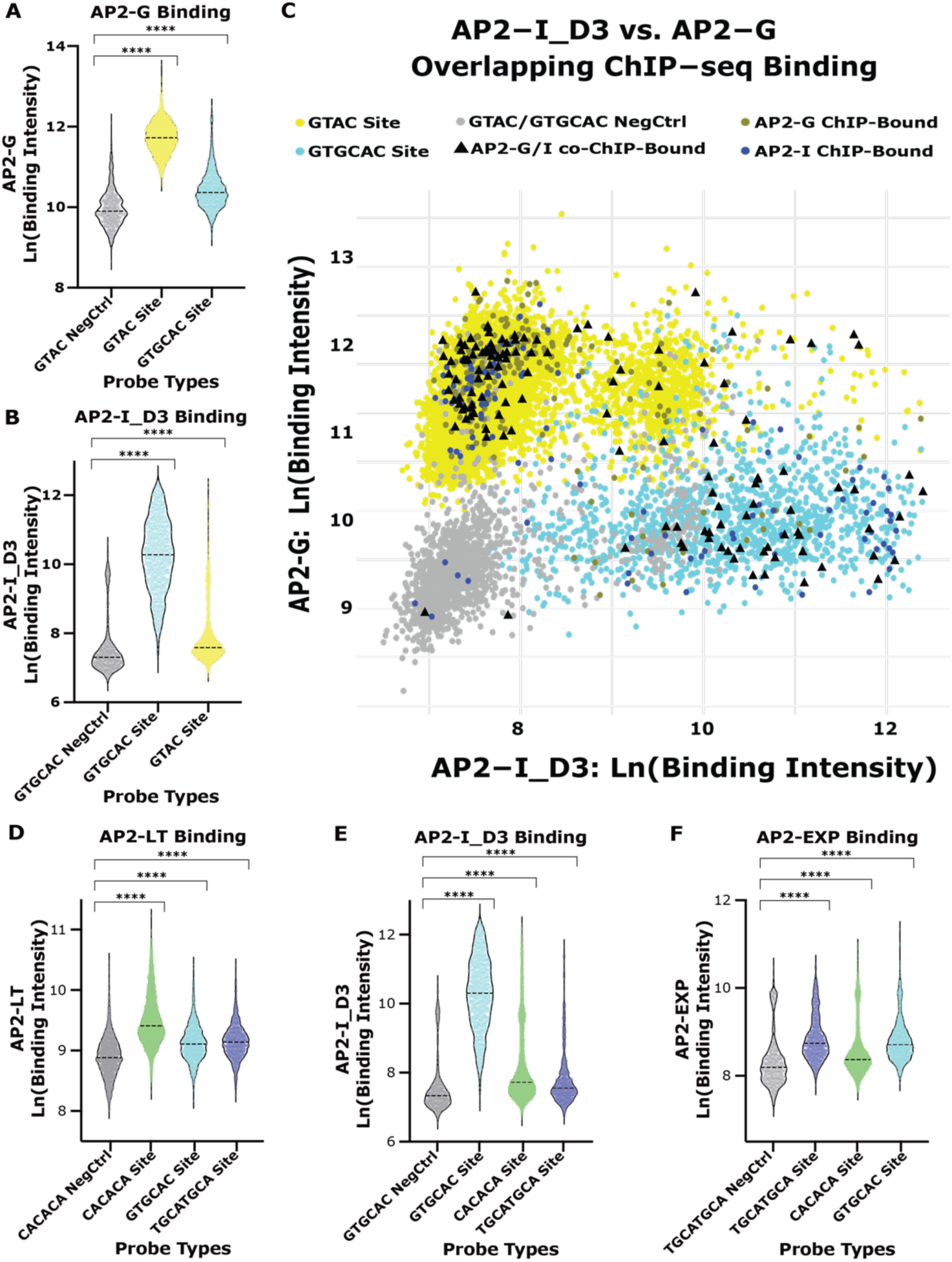
Overlapping *in vitro* binding preferences across DNA motif types (**A**) AP2-G binding intensity distributions for GTAC negative control probes (Grey), all GTAC probes (Yellow), and all GTGCAC probes (Blue); (**B**) AP2-I_D3 binding intensity distributions for GTGCAC negative control probes (Grey), all GTGCAC probes (Blue), and all GTAC probes (Yellow); (**C**) Comparison of the binding intensities for AP2-G and AP2-I_D3. Negative control probes (Grey), GTAC probes (Yellow), GTGCAC probes (Blue), AP2-G ChIP-bound sites (Dark Yellow), AP2-I ChIP-bound sites (Dark Blue), and AP2-G/AP2-I co-bound ChIP-bound sites (Black triangles); (**D**) AP2-LT binding intensity distributions for CACACA negative control probes (Grey), all CACACA probes (Green), all GTGCAC probes (Blue), and all TGCATGCA probes (Purple); (**E**) AP2-I_D3 binding intensity distributions for GTGCAC negative control probes (Grey), all GTGCAC probes (Blue), all CACACA probes (Green), and all TGCATGCA probes (Purple); (**F**) AP2-EXP binding intensity distributions for TGCATGCA negative control probes (Grey), all TGCATGCA probes (Purple), all GTGCAC probes (Blue), and all CACACA probes (Green).

Our AP2-LT ChIP-seq results identified an overlap with some AP2-EXP genome-wide binding sites, and previous work reported an overlap of AP2-I and AP2-EXP *in vivo* binding site preference^105^. Therefore, to expand the comparison to include all three factors, we identified the overlapping ChIP-seq peaks between AP2-LT (CACACA-binder), AP2-I (GTGCAC-binder), and AP2-EXP (TGCATGCA-binder) (**Supplemental Figure 7D**). Single-cell transcriptomics^74^ identified a subset of cells with co-expression of AP2-LT, AP2-I, and AP2-EXP (**Supplemental Figure 20K-N**), suggesting putative mechanisms of cooperative binding.

To characterize possible mechanisms of cooperative binding between AP2-LT, AP2-I, and AP2-EXP we first identified the genome-wide binding specificity for AP2-EXP using the *P. falciparum*-specific gcPBM. We found that AP2-EXP bound to TGCATGCA probes at higher signal intensities compared to the respective negative control probes with high correlation between technical replicates (**Supplemental Figure 23E-H**). AP2-EXP ChIP-bound sites had no significant difference compared to all TGCATGCA sites (**Supplemental Figure 15F**), and the *in vitro* versus *in vivo* binding comparison showed only a moderate correlation (**Supplemental Figure 24D**), which suggested no dependence on sequence context. AP2-EXP ChIP-bound sites were mostly defined by accessible chromatin and activating marks (**Supplemental Figure 15F**; **Supplemental Figure 25**).

We also analyzed the AP2-LT, AP2-I_D3, and AP2-EXP gcPBM data for binding to CACACA-, GTGCAC-, and TGCATGCA-containing probes. Each pairwise comparison (AP2- LT vs. AP2-I_D3, AP2-LT vs. AP2-EXP, and AP2-I_D3 vs. AP2-EXP) resulted in significant low affinity binding to probes containing other DNA motifs (**Figure 5D-F**), with a wide distribution of co-ChIP-bound sites (**Supplemental Figure 26**). Therefore, we find that some AP2 DBDs have the propensity to bind different DNA motifs *in vitro* at lower affinities which may drive the co-occupancy observed by *in vivo* ChIP-seq binding data.

## Discussion

Transcription factor (TF) binding site recognition and specificity are important components of gene regulation in all living organisms^1, 7, 8, 106–109^. Sequence-specific TFs classically recognize short, conserved DNA motifs. However, these specific DNA motifs may not fully capture biologically relevant binding site preferences at lower affinity genomic sites^28^. Moreover, features such as DNA sequence context, local DNA shape, protein-protein interaction partners, epigenetic post-translational modifications, and chromatin architecture also impact binding site recognition^5, 8, 11, 28, 106^. Investigating the parameters that differentiate the sequence specificity of large families of paralogous TFs in model eukaryotes poses a great challenge since they are often numerous (>100) and functionally redundant^9^. In this study, we explored the contribution of these context-dependent parameters in the malaria parasite *Plasmodium falciparum*, which has a reduced number of sequence-specific TFs relative to other eukaryotes ^5, 8, 11, 28, 110^

Using a novel *P. falciparum* genomic-context protein-binding DNA microarray (gcPBM), we found that three ApiAP2 TFs with DBDs that recognize a CACACA motif (PF3D7_0420300_D1, AP2-LT, and AP2-HC) cannot differentiate DNA sequence context *in vitro.* Despite this observation, we found that AP2-LT and AP2-HC^89^ do not bind overlapping genomic regions *in vivo.* We correlated our findings with previously published co-factor interactions, chromatin accessibility, and epigenetic mark datasets^67–69^, ^87–89, 119^ and found that chromatin and protein-complex features likely play important roles in defining the binding site selections *in vivo.* The absence of overlapping *in vivo* binding for AP2-LT and AP2-HC^89^ TFs is supported by recent studies that identified AP2-HC complexed with PfHP1^89^ at heterochromatic regions, while AP2-LT is a component of the putative PfSAGA transcriptional co-activating complex which is only found in euchromatic regions. While the specific role of PF3D7_0420300 as a TF during *P. falciparum* asexual blood stage development remains unknown, a previous publication identified its interaction with PfMORC^120^, a putative repressive complex component highly characterized in metazoans^121–128^ and recently in *Toxoplasma gondii*^129, 130^. Therefore, we anticipate that a PF3D7_0420300:PfMORC complex would bind genomic regions not bound by AP2-LT or AP2-HC. We also find that AP2-LT binds a double (TG/CA)CAC motif separated by five nucleotides *in vitro* and *in vivo*, suggesting a potential requirement for AP2-LT dimerization at genomic targets. Although a co-crystal structure of AP2-EXP:DNA revealed the possibility for AP2 domain-swapped homodimerization^62^, AP2-LT likely binds to the same face of the DNA via a different dimerization mechanism. In addition, we found that genomic sites bound by AP2-LT become accessible as the TF is being expressed at the mRNA^86^ and protein^98^ level, suggesting an interplay between TF occupancy and nucleosome positioning. Therefore, AP2-LT may recognize the TGCAC DNA motif on or near promoter-bound nucleosomes and recruit the PfSAGA chromatin remodeler complex^88^ to increase DNA accessibility. This potential activity for AP2-LT is reminiscent of pioneer factors in other eukaryotes^131–134^, but remains to be tested. We conclude that, while the three CACACA-binding TFs bound similar DNA sequence context specificities by gcPBM *in vitro*, most of the TFs had divergent chromatin preferences, suggesting that they are unlikely to be functionally redundant.

In contrast, our gcPBM results identified that SIP2_D1, AP2-I_D3, and HDP1, which bind the GTGCAC sequence, have differing *in vitro* preferences for nucleotides proximal to this core motif. These TFs also display a preference for divergent predicted DNA shape features, such as minor groove width and electrostatic potential, suggesting that shape-readout mechanisms^5^ impact binding site selection. Future structural work with these DBDs may help further define the base-and shape-readout mechanisms of these TF-DNA interactions. *In vivo* AP2-I prefers the AGTGCATTA extended motif and binds to regions of accessible chromatin containing activating epigenetic marks. SIP2 prefers a bipartite SPE2 GGTGCAC extended motif and colocalizes to sub-telomeric regions of inaccessible chromatin and repressive epigenetic marks^91^. For AP2-I and SIP2, a subset of cells from single-cell transcriptomics data^74^ show co-expression of both factors, suggesting the possibility of co-binding or competitive binding, which remains to be determined. Finally, HDP1 binds the extended TGTGCACA motif, yet is maximally expressed during sexual blood stages, for which comprehensive chromatin accessibility and epigenetic modifications datasets are lacking. Overall, we found that the GTGCAC-binding TFs have differential *in vitro* and *in vivo* DNA-binding preferences due to contributions from DNA sequence/shape context preferences, timing of expression, chromatin accessibility, and epigenetic patterns.

Our results also suggest that low-affinity protein-DNA interactions by *P. falciparum* DBDs may contribute to a more complex mechanism of *in vivo* TF co-occupancy. This concept has been investigated in the homeobox domain (HOX) TF family in *Drosophila*, where sites of Exd:Hox co-occupancies are driven by low-affinity binding between the TFs and DNA by latent specificity^28^. In *P. falciparum*, *in vivo* co-occupancy between AP2-G (GTAC-binder) and AP2-I (GTGCAC-binder) has been previously reported^83^. Here we further identified a subset of co-occupied sites by AP2-LT (CACACA-binder), AP2-I_D3^83^, and AP2-EXP^105^ (TGCATGCA- binder). It was previously hypothesized that, at sites of co-occupancy, one TF is driving the DNA-specific binding, while the other factor is present by protein-protein interactions^83^. However, using gcPBM binding data for the above DBDs, we found that each DBD bound divergent DNA motifs at low affinities, suggesting co-occupancy may be impacted by low-affinity binding from either factor. Overall, these results add to the growing evidence for genome-wide co-occupancy by ApiAP2 TFs.

This work contributes to our current understanding of how paralogous TF binding specificity is determined^45, 47, 56–58^. By interrogating sequence and chromatin features using *Plasmodium falciparum* TFs we characterized a reduced set of paralogous DNA binding domains from essential TFs with non-redundant functional roles. In this context, we found several solutions including TFs that rely on sequence context to differentiate genome-wide binding site selection and others that are coordinated through changes in chromatin state features. Our findings are also relevant from a therapeutic perspective since ApiAP2 TFs are unique to plant and eukaryotic parasite genomes and have therefore been proposed as future antiparasitic drug targets^105, 111–115^. Understanding the factors that drive complex gene regulatory mechanisms in *P. falciparum* is critical to selecting appropriate future drug interventions^116–118^. Recent work has identified putative antimalarial compounds that interact with AP2 domains *in silico* and *in vitro* which arrest *Plasmodium spp.* development at multiple stages of the parasite life cycle^105^. The concept of a “pan-ApiAP2 inhibitor” design would allow for the inhibition of multiple ApiAP2 TFs, such as the group of CACACA-binding TFs that regulate divergent developmental pathways, with one drug. Understanding the specific roles of these unique and parasite-essential factors will be critical for the design of future AP2-targeted antimalarial therapies.

## Acknowledgements

We would like to thank Riward Campelo Morillo and Björn F. C. Kafsack for providing us with recombinantly expressed and purified Homeodomain-like protein 1 (HPD1) samples, Timothy J. Russell and Gabriel W. Rangel for proof-reading this manuscript, and Lindsey M. Orchard for technical assistance.

## Author Contributions

Victoria A. Bonnell: project conception, data generation, data analysis, manuscript writing. Yuning Zhang: data analysis, manuscript editing. Alan S. Brown: data analysis, manuscript editing. John Horton: data generation. Gabrielle A. Josling: data generation. Tsu-Pei Chiu: data analysis, manuscript editing. Remo Rohs: manuscript editing. Shaun Mahony: data analysis, manuscript editing. Raluca Gordân: data analysis, manuscript editing. Manuel Llinás: project conception, data analysis, manuscript writing.

