## Supplemental Figures for "DNA sequence context and the chromatin landscape differentiate sequence-specific transcription factor binding in the human malaria parasite, *Plasmodium falciparum*"

### Supplementary Figures:

**A**

| Gene Name | AA Start | AA End |
|---|---|---|
| PF3D7_0423000_D1 | 1956 | 2015 |
| PF3D7_0820100_D1 (AP2-LT_D1) | 2520 | 2577 |
| PF3D7_1456000_D1 (AP2-HC_D1) | 1316 | 1374 |
| PF3D7_0604100_D1 (SIP2_D1) | 177 | 235 |
| PF3D7_1007700_D3ext (AP2-I_D3) | 910 | 1027 |
| PF3D7_1466200_D1 (HDP1_D1) | 2990 | 3079 |
| PF3D7_1222600_D1ext (AP2-G_D1) | 2123 | 2245 |
| PF3D7_1466400_D1 (AP2-EXP_D1) | 63 | 123 |

**B**

```
PF3D7_0420300_D1    GPLPTGVYFDSARKLWRCQWKENGKFKTKGFSLIHYSTLEEARKQCILYRCDVGNIPVKS
AP2LT_D1            GEKVSGVWYDTNRRLWRVMYMENDKRKTRGFSPRIYG-FNAARDLAVQLKYEMNKKNK-K
AP2HC_D1            FEARKGVYYDKSRKLWRANWKENGKIQTKGFSVNEYKSVQLARQKAIEWREKKEAELL-L
Top 3 Consensus         .**::*. *:***  :  **.*  :*:***   *  .:  **.  .:  :  .
                                              ■   ■   ■
PF3D7_1305200_D1    KSGYRGVFYEPSYHAYICIYNEAGVRKFQIFKIQNNDY-LEAYHLALMCRRYKLFKNYQF
All 4 Consensus          **:::   : :   :  * .  : :  *.         * . .:  :
```

| CACACA | PF3D7_0420300_D1 | PF3D7_0802100_D1 (AP2-LT_D1) | PF3D7_1305200_D1 | PF3D7_1456000_D1 (AP2-HC_D1) | PF3D7_1466400_D1 (AP2-EXP_D1) |
|---|---|---|---|---|---|
| PF3D7_0420300_D1 | 100 | 33.33 (48.15) | 19.61 (41.18) | 49.02 (62.75) | 14.81 (37.04) |
| PF3D7_0802100_D1 (AP2-LT_D1) | - | 100 | 22.64 (39.62) | 35.19 (55.58) | 20.37 (46.30) |
| PF3D7_1305200_D1 | - | - | 100 | 18.87 (41.51) | 16.67 (42.59) |
| PF3D7_1456000_D1 (AP2-HC_D1) | - | - | - | 100 | 20.37 (42.59) |
| PF3D7_1466400_D1 (AP2-EXP-D1) | - | - | - | - | 100 |

**C**

```
SIP2_D1      IGSQEPVILIDKIERCLVVEWYENNIRREQRIS---YKKYGNDKAKLRAKELIEKLKSGITF-
AP2I_D3      LTPVEGV-RFDPYSYSWFAKYLENENSKEPKISKYLLKKWGFNKAHSLAVHTVKCAYKAVPFT

Consensus    : . * *  :*  . . ..:: **:  :* :**    **:* :**:  * . ::   ..:.*
```

| GTGCAC | PF3D7_0604100_D1 (SIP2_D1) | PF3D7_1007700_D3 (AP2-I_D3) | PF3D7_1466400_D1 (AP2-EXP_D1) |
|---|---|---|---|
| PF3D7_0604100_D1 (SIP2_D1) | 100 | 21.88 (32.81) | 15.52 (32.76) |
| PF3D7_1007700_D3 (AP2-I_D3) | - | 100 | 16.92 (36.92) |
| PF3D7_1466400_D1 (AP2-EXP_D1) | - | - | 100 |

**Supplemental Figure 1: Amino acid sequence alignments of DBDs used in this study**

**(A)** Amino acid coordinates of DBDs used for gcPBM experiments as determined previously[42,43,48]; **(B)** *Above:* Amino acid sequence alignment of CACACA-binding AP2 domains[110]. *Below:* Calculated percent identity (and percent similarity) from the CACACA-binding alignment[111], with AP2-EXP as an outgroup; **(C)** *Above:* Amino acid sequence alignment of GTGCAC-binding AP2 domains[110]. *Below:* Calculated percent identity (and percent similarity) from the GTGCAC-binding alignment[111], with AP2-EXP as an outgroup.

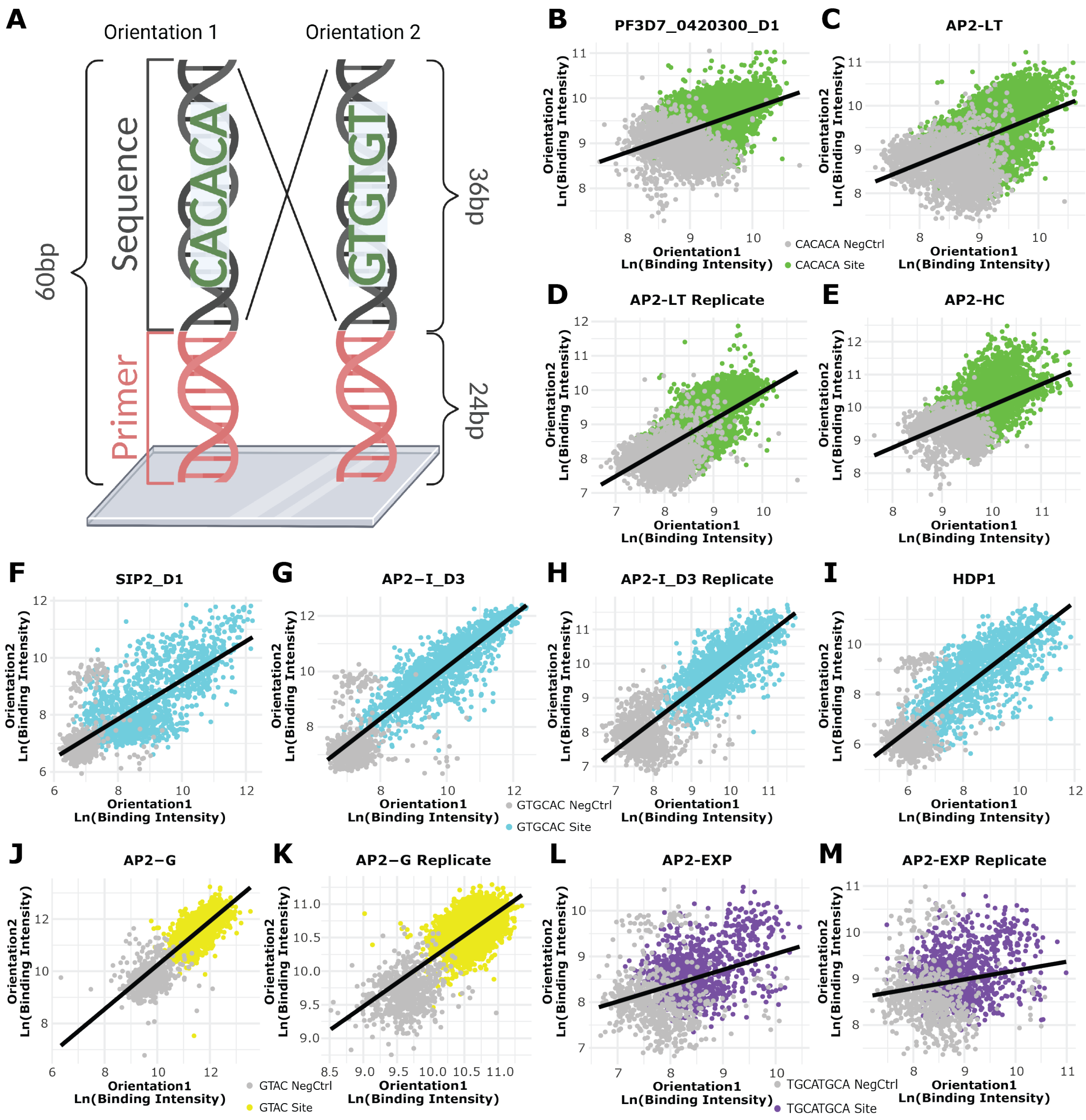


**Supplemental Figure 2: Differences in DNA-binding across probe orientations**

**(A)** Graphical representation of the two orientations from gcPBM experiments; **(B-E)** Comparison of binding intensity across probe orientations for the CACACA-binding group including the technical replicate (CACACA probes [Green] and negative control probes [Grey]). Line of best fit the GTGCAC-binding group including the technical replicate (GTGCAC probes [Blue] and negative control probes [Grey]); **(J-K)** Comparison of binding intensity across probe orientations for AP2-G including the technical replicate (GTAC probes [Yellow] and negative control probes [Grey]); **(L-M)** Comparison of binding intensity across probe orientations for AP2-EXP including a technical replicate (TGCATGCA probes [Purple] and negative control probes [Grey]). Microarray graphic generated using BioRender..

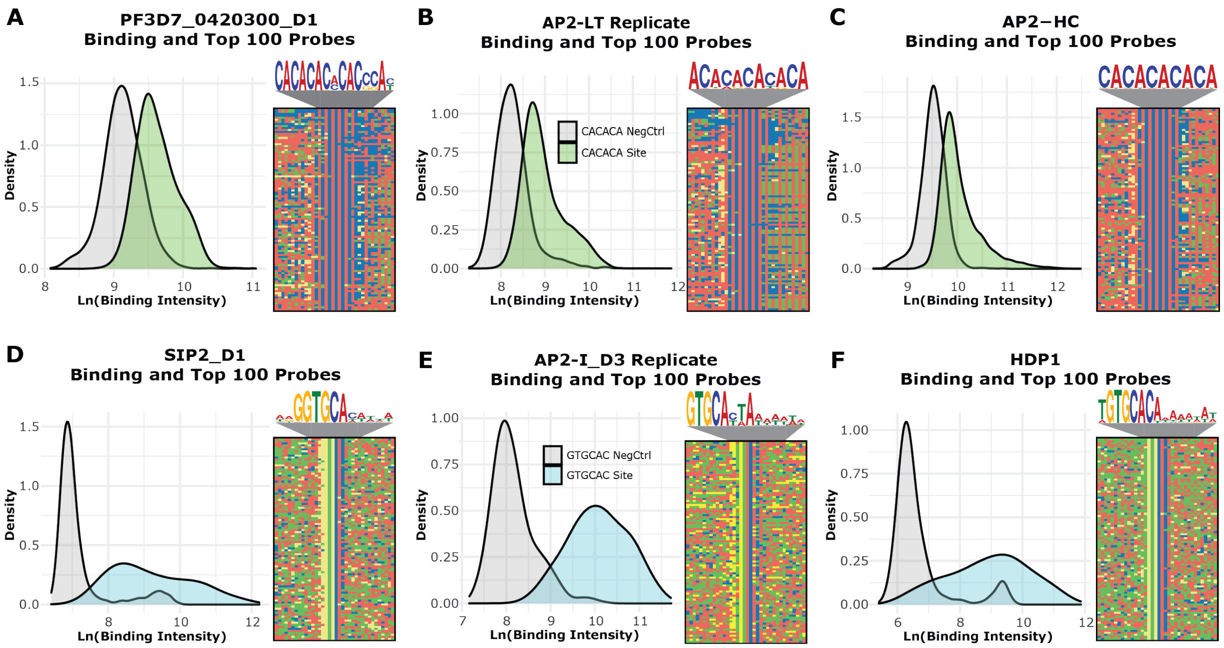


**<u>Supplemental Figure 3: Distribution of binding specificity and top 100 bound probes for DBDs</u>**

**(A-C)** *Left*: Distributions of the binding intensities for the CACACA-binding group including the technical replicate (CACACA probes [Green] and negative control probes [Grey]). *Right*: the DNA motif enriched[86] in the 100 top bound probes with a four-color plot of the 100 top bound probes underneath. Color representations: A (Red), C (Blue), G (Yellow), and T (Green); **(D-F)** *Left*: Distributions of the binding intensities for the GTGCAC-binding group including a technical replicate (GTGCAC probes [Blue] and negative control probes [Grey]).

**A**

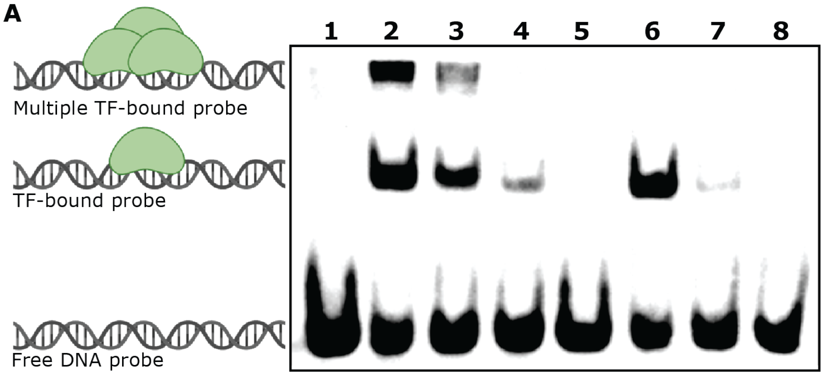


| Lane # | DNA Sequence | DBD Added |
|---|---|---|
| 1 | CATATTCCCCCCACACACACACACGCATACATACC | No Protein |
| 2 | CATATTCCCCCCACACACACACACGCATACATACC | AP2-LT |
| 3 | CATATTCCCCCCTTTTACACACACGCATACATACC | AP2-LT |
| 4 | CATATTCCCCCCACACTTTTACACGCATACATACC | AP2-LT |
| 5 | CATATTCCCCCCACACACACACACGCATACATACC | No Protein |
| 6 | CATATTCCCCCCACACACACACACGCATACATACC | AP2-HC |
| 7 | CATATTCCCCCCTTTTACACACACGCATACATACC | AP2-HC |
| 8 | CATATTCCCCCCACACTTTTACACGCATACATACC | AP2-HC |

**B**

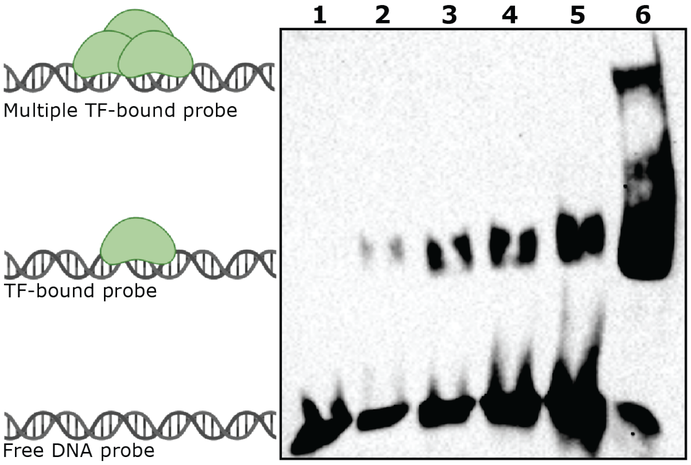


| Lane # | DNA Sequence | DBD Added | Ln(Binding Intensity) |
|---|---|---|---|
| 1 | TGAAAAAAAAAAAAAAAAAAAAGAAAAAAAAAAAAA | AP2-LT | 8.213 |
| 2 | TTTTTTTTTCATTCACACAAAAAAAAAAAATTTAT | AP2-LT | 8.305 |
| 3 | TTTTTGTATTCTCCGGACTACATTATGTTGTAAATG | AP2-LT | 9.467 |
| 4 | TAAAATATAATGATCACACATATAAACAAGTGGGTA | AP2-LT | 9.255 |
| 5 | AATAATAGAAGCACCCACCCACCACCACCCACACAT | AP2-LT | 10.479 |
| 6 | CATATTCCCCCCACACACACACACGCATACATACC | AP2-LT | 10.985 |

**Supplemental Figure 4: AP2-LT DBD binds to longer CA-dinucleotides with multiple DBDs** **(A)** Electrophoretic mobility shift assays (EMSAs) with purified AP2-LT and AP2-HC AP2 domains. The DNA sequences associated with the lane numbers are below the gel image (ChemiDoc exposure time 25sec). The protein added to lanes noted below the gel image. The "No protein" notation represents only free probe without protein added as a negative control; **(B)** Validation EMSA with purified AP2-LT AP2 domain with different DNA probes that had low-to-high binding in the gcPBM experiments (*from left to right*). (ChemiDoc exposure time 600sec) Protein-DNA interaction graphic generated using BioRender.

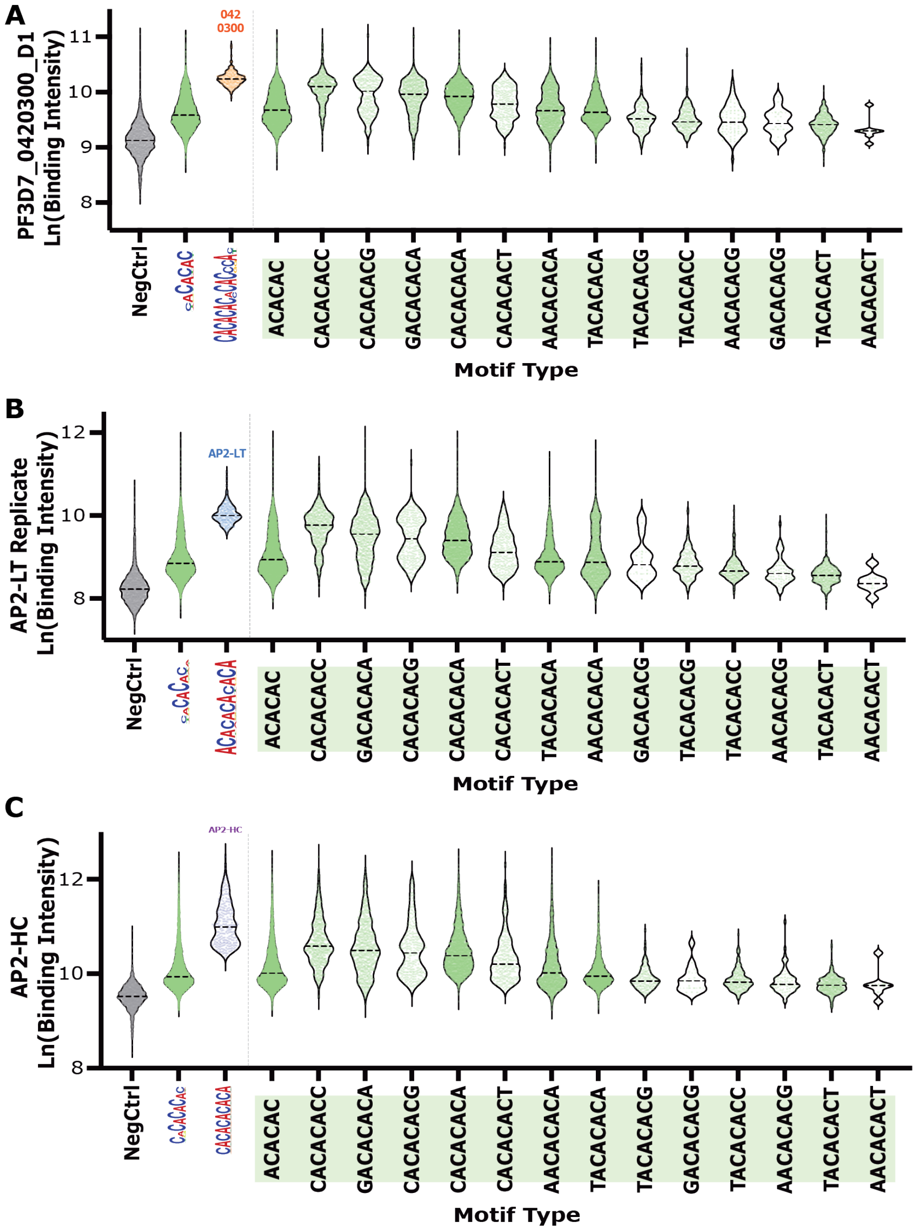


**Supplemental Figure 5: CACACA-binding DBDs have minimal differences in sequence context preferences**

**(A)** Binding intensity distributions for CACACA negative control probes (Grey), all CACACA probes (Green), the PF3D7_0420300_D1 extended motif probes (Orange), and 8-mer CACACA probes represented in the gcPBM (Green). Dotted lines are the calculated mean for each violin plot; **(B)** Binding intensity distributions for CACACA negative control probes (Grey), all CACACA probes (Green), the AP2-LT Replicate extended motif probes (Blue), and 8-mer CACACA probes represented in the gcPBM (Green). Dotted lines are the calculated mean for each violin plot; **(C)** Binding intensity distributions for CACACA negative control probes (Grey), all CACACA probes (Green), the AP2-HC extended motif probes (Purple), and 8-mer CACACA probes represented in the gcPBM (Green). Dotted lines are the calculated mean for each violin plot;

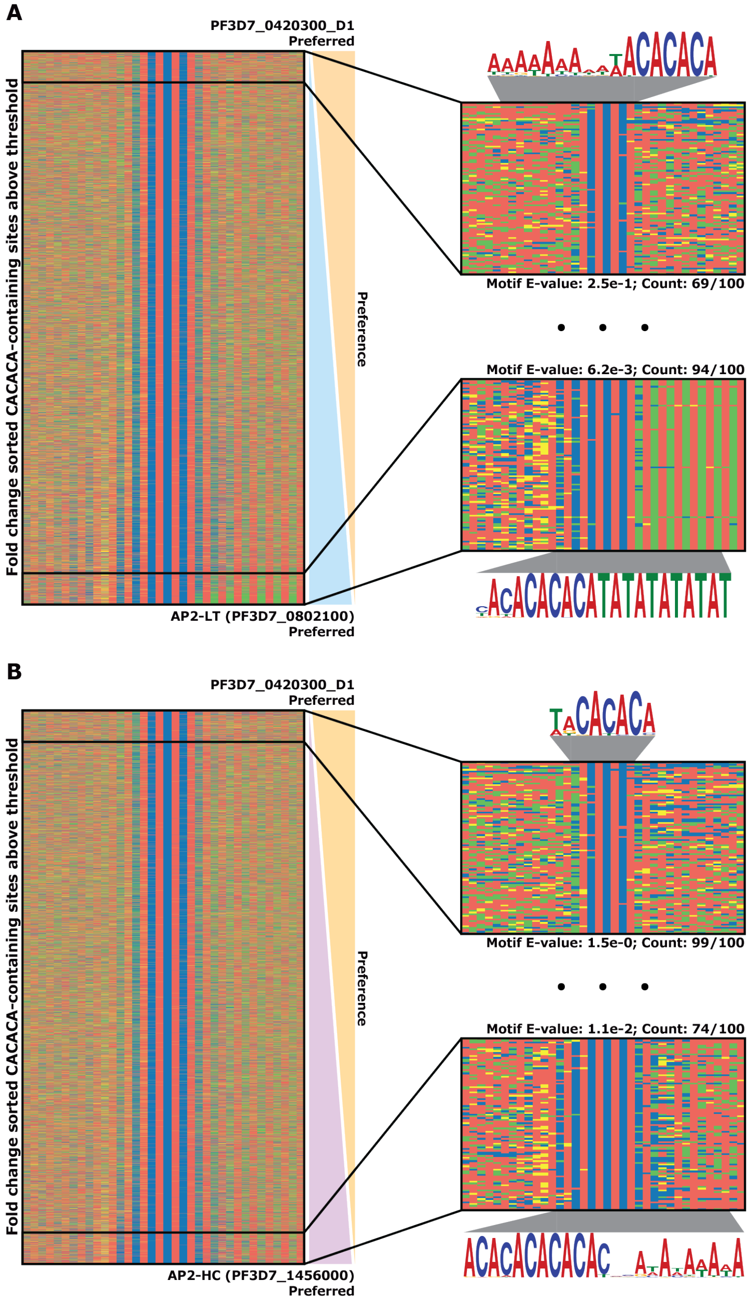


**Supplemental Figure 6: Differential sequence preferences of CACACA-binding DBDs**

***(A)*** *Left:* Four-color plot of CACACA probes above the 90th percentile of negative control probes sorted by fold change ($\log_2$[PF3D7_0420300/AP2-LT]). *Right:* Zoom in on the top 100 differentially bound probes by PF3D7_0420300_D1 (*top right*) and AP2-LT (*bottom right*) with enriched motifs, calculated E-values, and motif occurrence counts within the top 100 sites[86]. Color representations: A (Red), C (Blue), G (Yellow), and T (Green); **(B)** *Left:* Four-color plot of CACACA probes above the 90th percentile of negative control probes sorted by fold change ($\log_2$[PF3D7_0420300/AP2-HC]). *Right:* Zoom in on the top 100 differentially bound probes by PF3D7_0420300_D1 (*top right*) and AP2-HC (*bottom right*) with enriched motifs, calculated E-values, and motif occurrence counts within the top 100 sites[86].

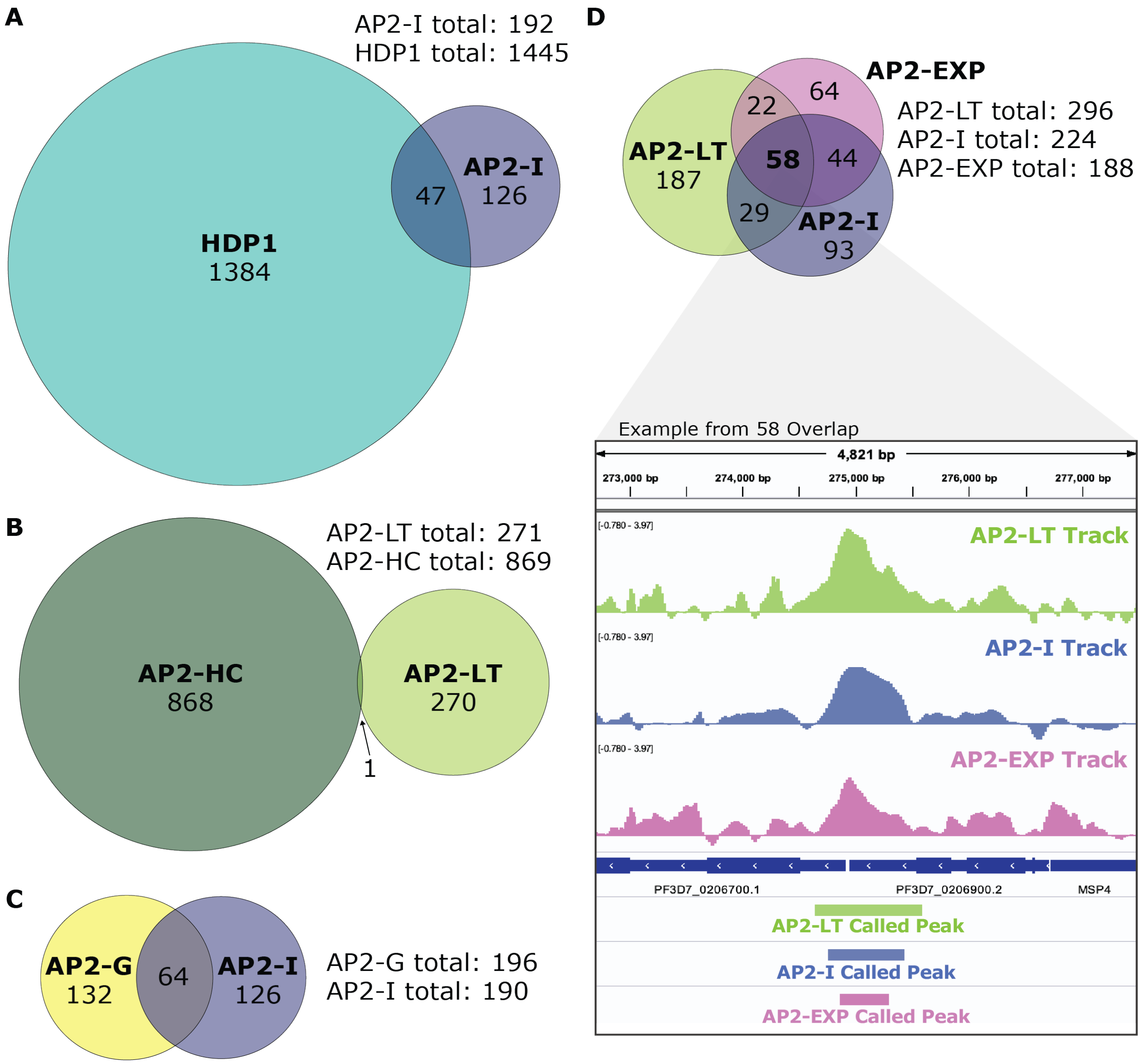


**Supplemental Figure 7: Comparisons of *in vivo* genome-wide occupancies of *P. falciparum* TFs in this study**

**(A)** Comparison of the genomic regions bound by HDP1 and AP2-I from published ChIP-seq experiments[48,84,90,95]; (**B**) Comparison of the genomic regions bound by AP2-LT (this study) and published AP2-HC; (**C**) Comparison of the genomic regions bound by published AP2-G and AP2-I; (**D**) Comparison of the genomic regions bound by AP2-LT (this study), published AP2-I, and published AP2-EXP. *Bottom*: screenshot from Integrative Genomics Viewer (IGV)[107] of a representative overlapping locus with AP2-LT (Green), AP2-I (Blue), and AP2-EXP (Pink). Significantly bound regions identified by MACS2 peak caller[85,99] represented as genomic interval bars under peak data.

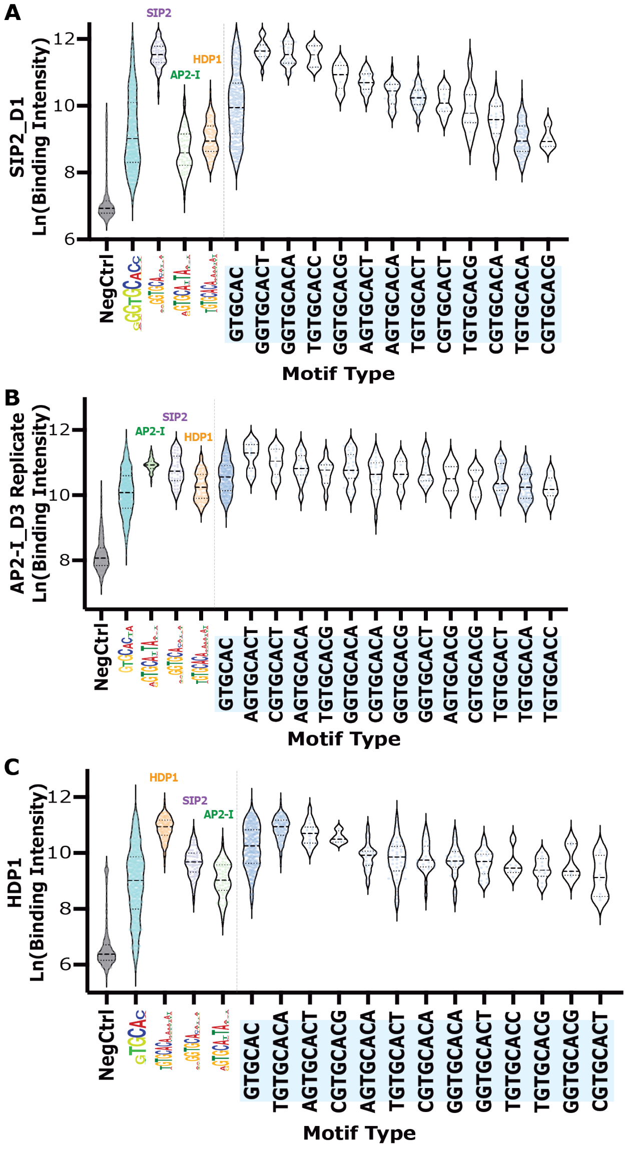


**Supplemental Figure 8: GTGCAC-binding DBDs have divergent sequence context preferences**

**(A)** Binding intensity distributions of SIP2_D1 for GTGCAC negative control probes (Grey), all GTGCAC probes (Blue), the extended motif probes by all three GTGCAC-binding TFs (AP2-I_D3[Green], SIP2_D1[Purple], and HDP1[Orange]), and 8-mer GTGCAC probes represented in the gcPBM (Blue). Dotted lines are the calculated mean for each violin plot; **(B)** Binding intensity distributions of AP2-I_D3 Replicate for GTGCAC negative control probes (Grey), all GTGCAC probes (Blue), the extended motif probes by all three GTGCAC-binding TFs (AP2-I_D3[Green], SIP2_D1[Purple], and HDP1[Orange]), and 8-mer GTGCAC probes represented in the gcPBM (Blue). Dotted lines are the calculated mean for each violin plot; **(C)** Binding intensity distributions of HDP1 for GTGCAC negative control probes (Grey), all GTGCAC probes (Blue), the extended motif probes by all three GTGCAC-binding TFs (AP2-I_D3[Green], SIP2_D1[Purple], and HDP1[Orange]), and 8-mer GTGCAC probes represented in the gcPBM (Blue). Dotted lines are the calculated mean for each violin plot.

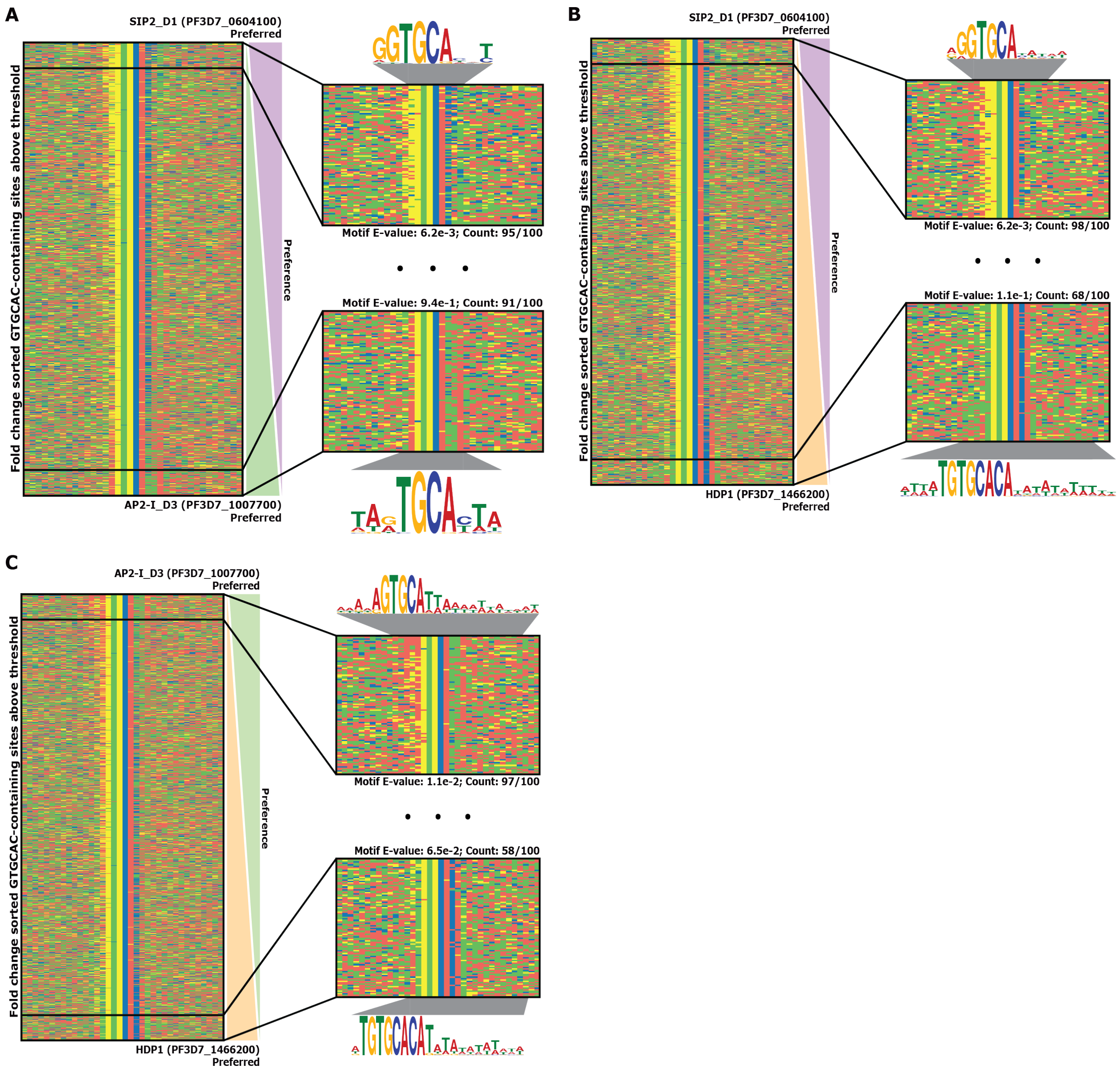


**Supplemental Figure 9: Differential sequence preference of GTGCAC-binding TFs**

**(A)** *Left*: Four-color plot of all GTGCAC probes in gcPBM design above 90$^{th}$ percentile of the negative control probes, sorted by fold change (Log$_2$[SIP2/AP2-I]). *Right*: Enriched DNA motif[86] and zoom in of the top 100 bound probes by each SIP2_D1 (*top*) and AP2-I_D3 (*bottom*). Color representations: A (Red), C (Blue), G (Yellow), and T (Green); (**B**) *Left*: Four-color plot of all GTGCAC probes in gcPBM design above 90$^{th}$ percentile of the negative control probes, sorted by fold change (Log$_2$[SIP2/HDP1]). *Right*: Enriched DNA motif[86] and zoom in of the top 100 bound probes by each SIP2_D1 (*top*) and HDP1 (*bottom*); (**C**) *Left*: Four-color plot of all GTGCAC probes in gcPBM design sorted by fold change (Log$_2$[AP2-I/HDP1]). *Right*: Enriched DNA motif[86] and zoom in of the top 100 bound probes by each AP2-I_D3 (*top*) and HDP1 (*bottom*).

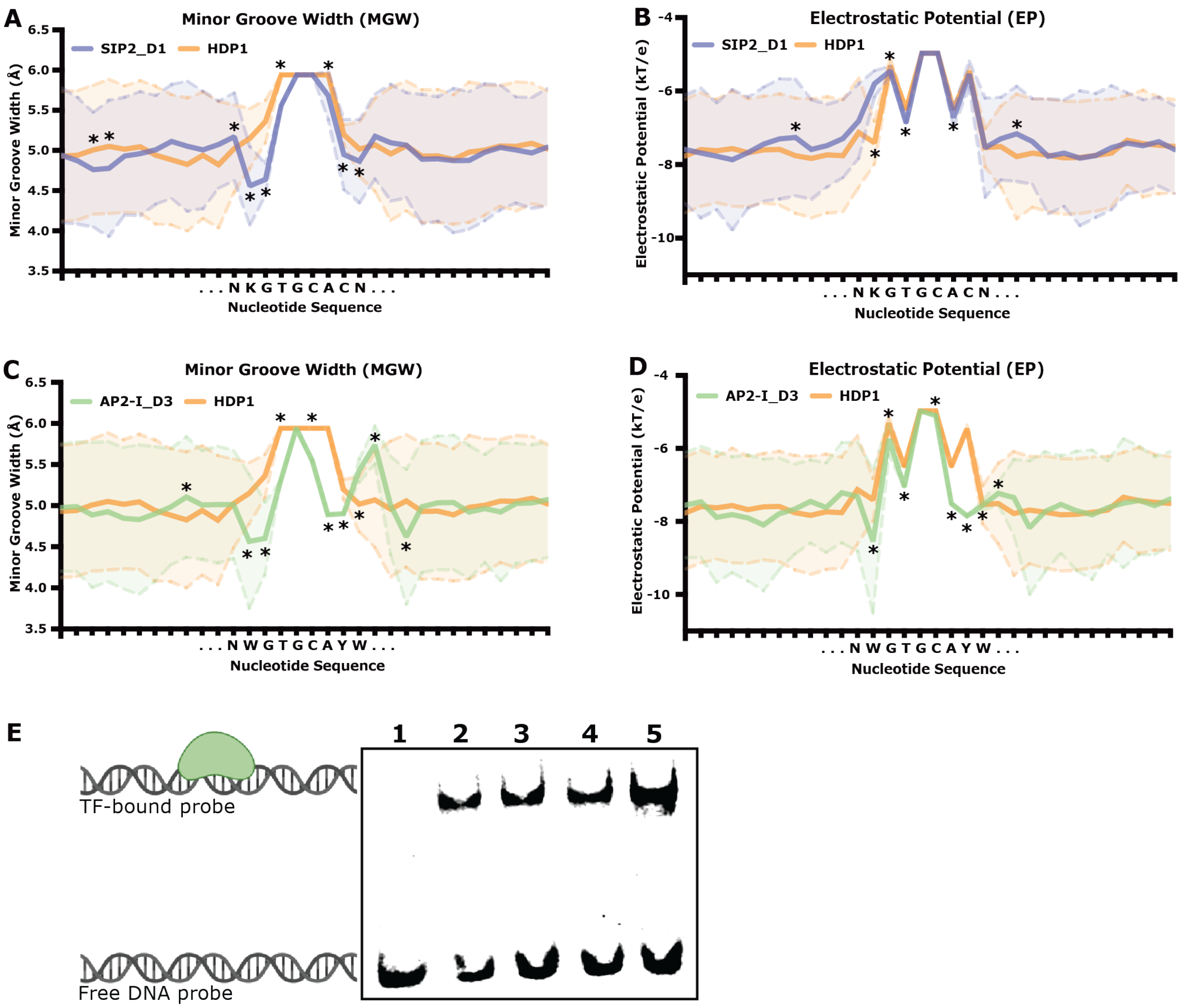


| Lane # | DNA Sequence | DBD Added |
|---|---|---|
| 1 | GATAAAATGAGACCAGTGCATTATTAGTTGTACACC | No Protein |
| 2 | GATAAAATGAGACCAGTGCATTATTAGTTGTACACC | AP2-I_D3 |
| 3 | GATAAAATGAGACCAGTGCATTATTTGTTGTACACC | AP2-I_D3 |
| 4 | GATAAAATGAGAAAAGTGCATTATTTGTTGTACACC | AP2-I_D3 |
| 5 | GATAAAATGAGTAAAGTGCATTAAATGTTGTACACC | AP2-I_D3 |

**Supplemental Figure 10: Differential DNA shape preference of GTGCAC-binding TFs**
**(A-B)** Calculated minor groove width (MGW) and electrostatic potential (EP) profiles[83] for SIP2_D1-preferred-GGTGCAC probes (Purple) and HDP1-preferred-TGTGCACA (Orange) probes. Significant differences (p-value < 0.05) between features notated with an * (two-sided Wilcoxon rank sum test). N = IUPAC for any nucleotide. K = IUPAC for G or T nucleotides; **(C-D)** Calculated MGW and EP profiles[83] for AP2-I_D3-preferred-AGTGCATTA (Green) probes and HDP1-preferred-TGTGCACA (Orange) probes. Significant differences (p-value < 0.05) between features notated with an * (two-sided Wilcoxon rank sum test). N = IUPAC for any nucleotide. W = IUPAC for A or T nucleotides. Y = IUPAC for C or T nucleotides; (**E**) EMSA of AP2-I_D3 binding to a AGTGCATTA probes with increasing numbers of shape mutants to the sequences outside the extended motif. Protein-DNA interaction graphic generated using BioRender.

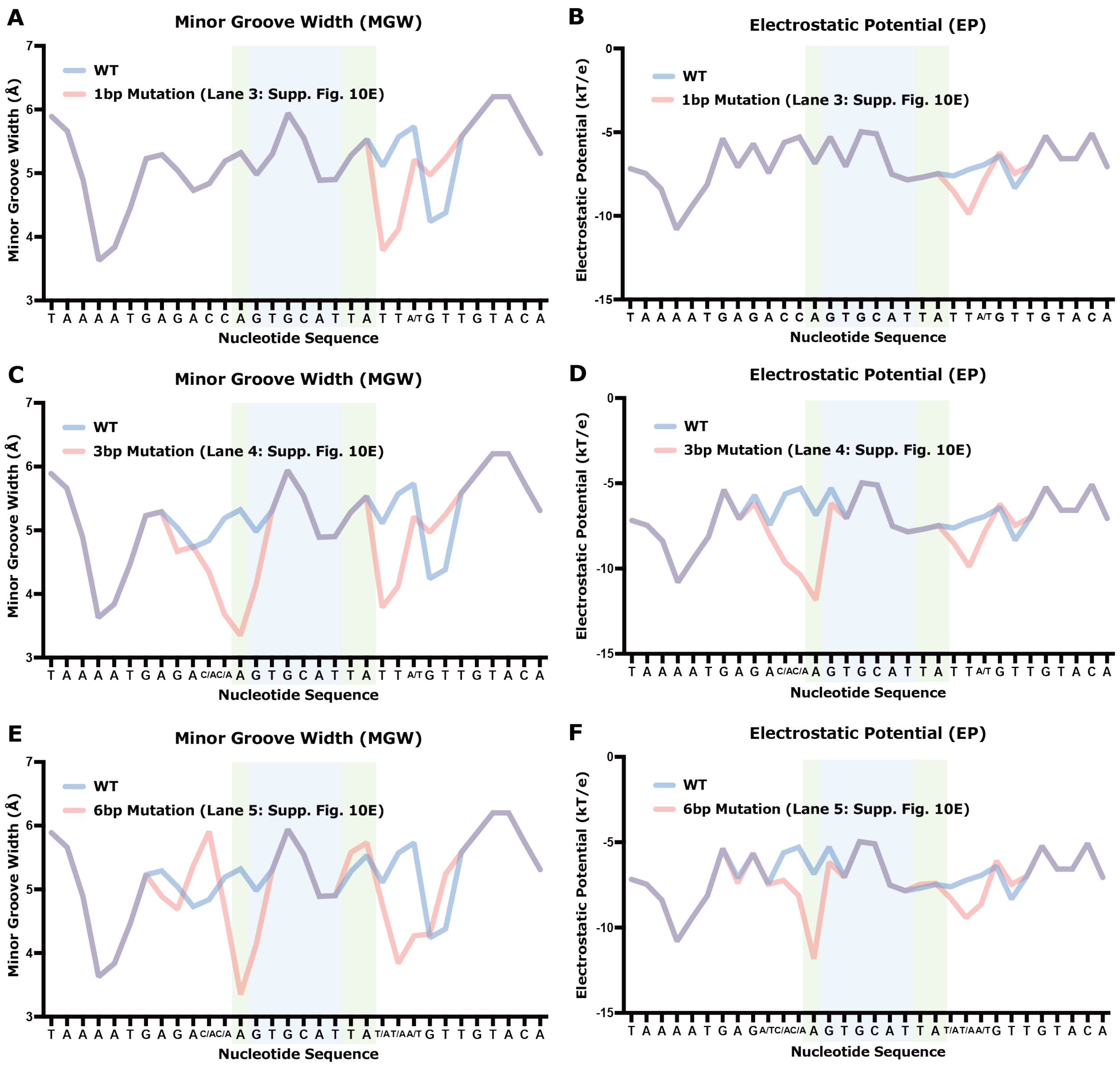


**<u>Supplemental Figure 11: Modeled DNA shape profiles of distal shape mutations</u>**

Calculated minor groove width (MGW) and electrostatic potential (EP) DNA shape profiles[83] of the WT (Blue) and mutated (Pink) DNA probes for electrophoretic mobility shift assays (EMSAs). Blue window highlights the GTGCAC sequence and the Green window highlights the extended motif proximal to the GTGCAC sequence on each figure. (**A-B**) MGW and EP profiles of one point mutation from **Supplemental Figure 10E**; (**C-D**) MGW and profiles of three point mutations from **Supplemental Figure 10E**; (**E-F**) MGW and EP profiles of six point mutations from **Supplemental Figure 10E**.

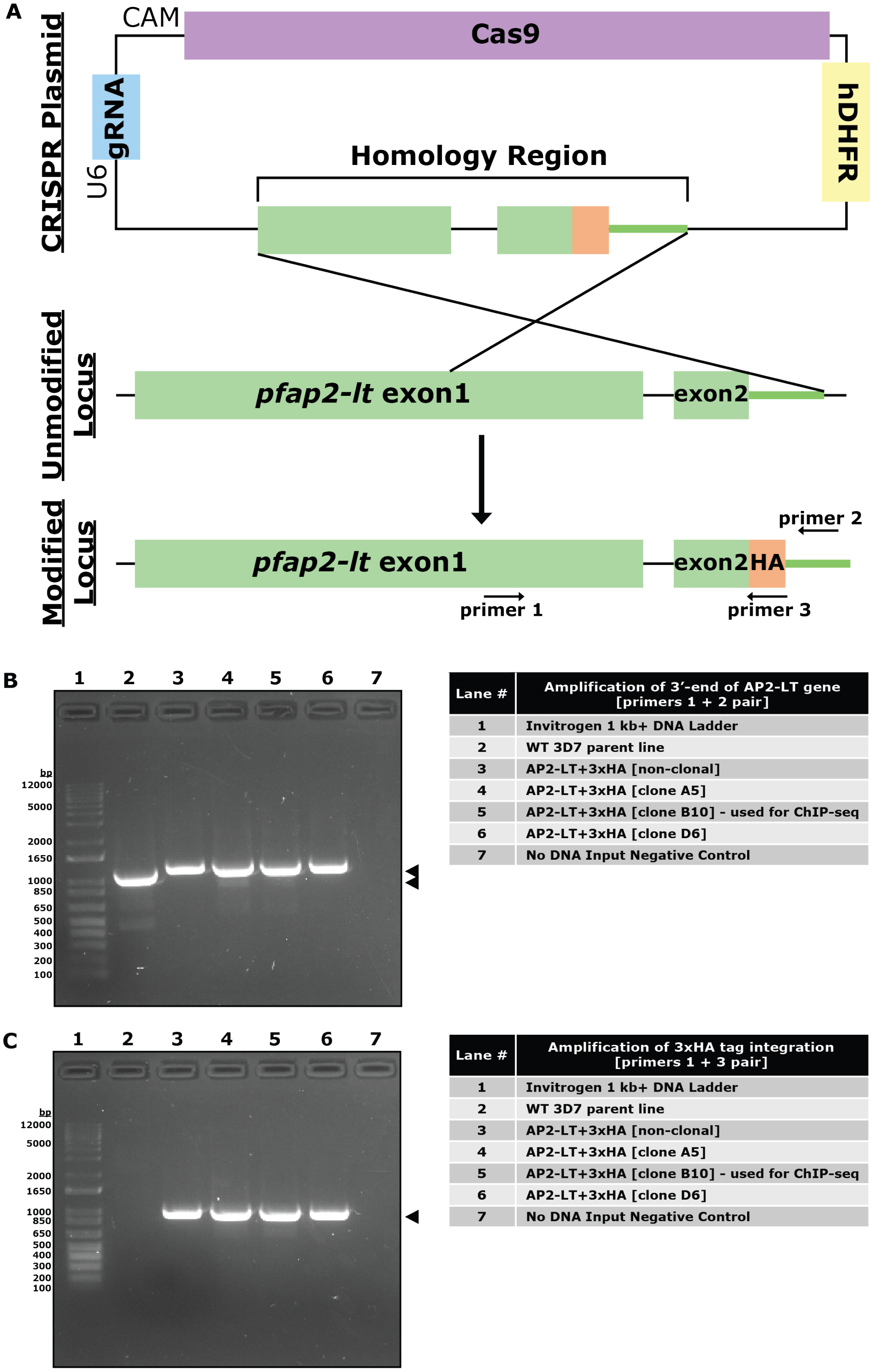


| Lane # | Amplification of 3'-end of AP2-LT gene [primers 1 + 2 pair] |
|---|---|
| 1 | Invitrogen 1 kb+ DNA Ladder |
| 2 | WT 3D7 parent line |
| 3 | AP2-LT+3xHA [non-clonal] |
| 4 | AP2-LT+3xHA [clone A5] |
| 5 | AP2-LT+3xHA [clone B10] - used for ChIP-seq |
| 6 | AP2-LT+3xHA [clone D6] |
| 7 | No DNA Input Negative Control |

| Lane # | Amplification of 3xHA tag integration [primers 1 + 3 pair] |
|---|---|
| 1 | Invitrogen 1 kb+ DNA Ladder |
| 2 | WT 3D7 parent line |
| 3 | AP2-LT+3xHA [non-clonal] |
| 4 | AP2-LT+3xHA [clone A5] |
| 5 | AP2-LT+3xHA [clone B10] - used for ChIP-seq |
| 6 | AP2-LT+3xHA [clone D6] |
| 7 | No DNA Input Negative Control |

**Supplemental Figure 12: Generation of PfAP2-LT$^{HA}$ tagged line for ChIP-seq experiments**
**(A)** Schematic of endogenous gene locus tagging by double homologous recombination via CRISPR-Cas9; **(B)** Genotyping PCR to amplify the 3`-end of *pfap2-lt* with primers 1 and 2 (notated in Panel A); **(C)** Genotyping PCR to amplify only if the tag was integrated into the endogenous *pfap2-lt* locus with primers 1 and 3 (notated in Panel A).

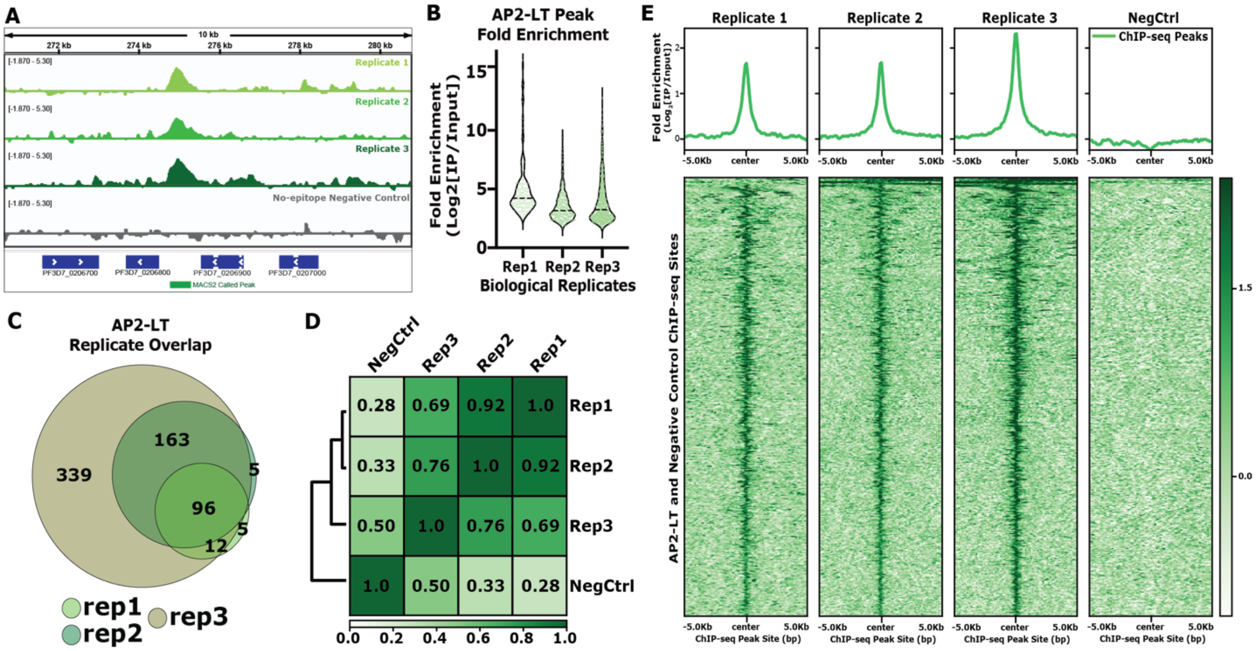


**Supplemental Figure 13: Quality control for AP2-LT ChIP-seq experiments**

(**A**) Example screenshot from Integrative Genomics Viewer (IGV)[107] of a 10kb region on chromosome 2. ChIP-seq genome tracks depicted from top to bottom are as follows: AP2-LT biological replicate one ($log_2$[IP/Input]) [Light Green], AP2-LT biological replicate two ($log_2$[IP/Input]) [Medium Green], AP2-LT biological replicate three ($log_2$[IP/Input]) [Dark Green], No-epitope Negative Control single replicate ($log_2$[IP/Input]) [Dark Grey], *P. falciparum* 3D7 strain gene annotation (Pfalciparum3D7, version 3, release 38[81]) [Dark Blue], and significantly (q-value < 0.01) called peak region by MACS2[85,99] [Green bar]; (**B**) Distribution of fold enrichment ($Log_2$[IP/Input]) of AP2-LT ChIP-seq MACS2-called peaks from each replicate experiment; (**C**) MACS2-called peak ChIP-seq replicate overlaps; (**D**) Pearson correlation plot of AP2-LT ChIP-seq three biological replicates and no-epitope negative control ChIP-seq sample; (**E**) *Top:* Profile plot of the mean AP2-LT ChIP-seq fold enrichment ($Log_2$[IP/Input]) across all three biological replicates and no-epitope negative control sample. *Bottom:* Heatmap of the AP2-LT ChIP-seq fold enrichment ($Log_2$[IP/Input]) across all three biological replicates and no-epitope negative control sample.

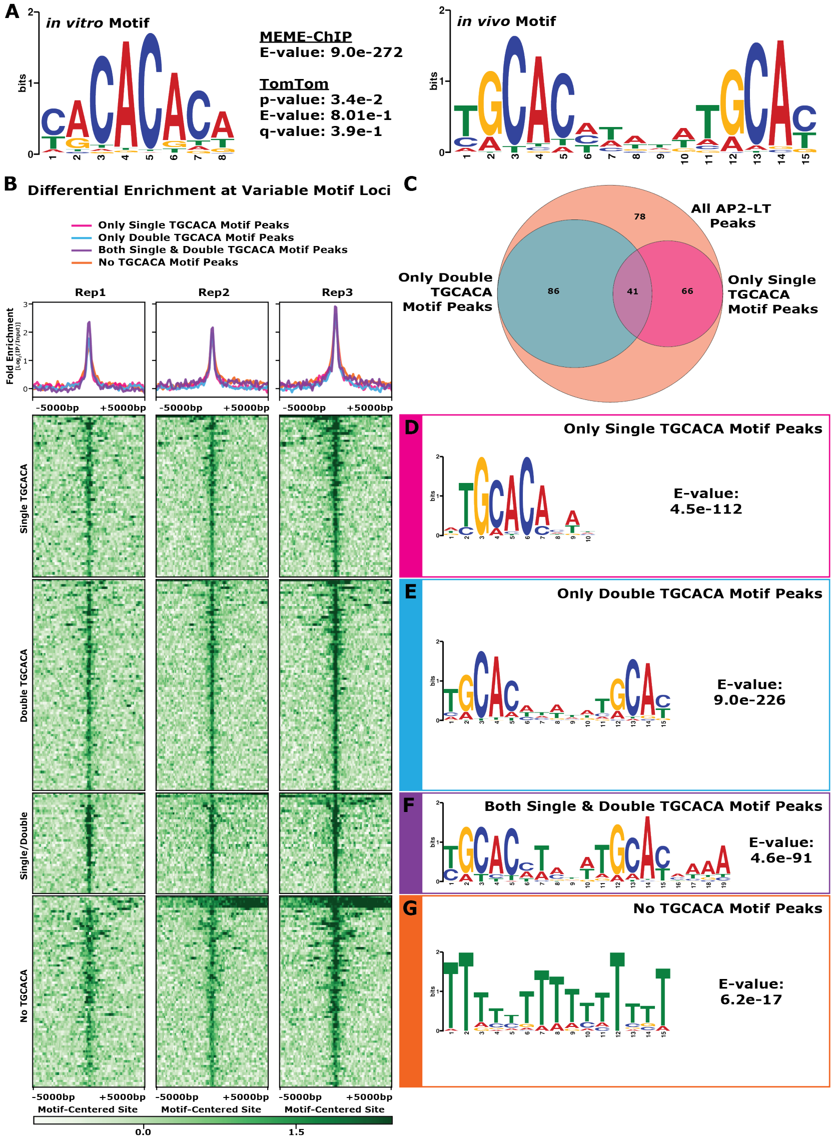


**Supplemental Figure 14: Motif enrichment analyses for single and double TGCAC motif in AP2-LT ChIP-seq binding sites**

(**A**) Enriched CACACA motif from previous *in vitro* universal Protein-Binding Microarray[43] and TGCACN$_5$TGCAC enriched DNA motif *in vivo* (this study), calculated motif E-value for the double motif via MEME-ChIP analysis, and p-, E-, and q-values for motif comparison via TomTom analysis[86]; (**B**) *Top:* Profile plot of the mean AP2-LT ChIP-seq fold enrichment (Log$_2$[IP/Input]) across all three biological replicates and categorized into four different regions (Only single TGCAC motif peaks [Pink], only double TGCAC motif peaks [Blue], both single and double motif peaks [Purple], and no TGCAC motif peaks [Orange]). *Bottom:* Heatmap of the AP2-LT ChIP-seq fold enrichment (Log$_2$[IP/Input]) across all three biological replicates and categorized into four different regions (shown above); (**C**) Overlap of the four region categories; **(D)** Motif enrichment and E-value for only single TGCAC motif peaks; **(E)** Motif enrichment and E-value for only double TGCAC motif peaks; **(F)** Motif enrichment and E-value for both single and double TGCAC motif peaks; and **(G)** Motif enrichment and E-value for No TGCAC motif peaks.

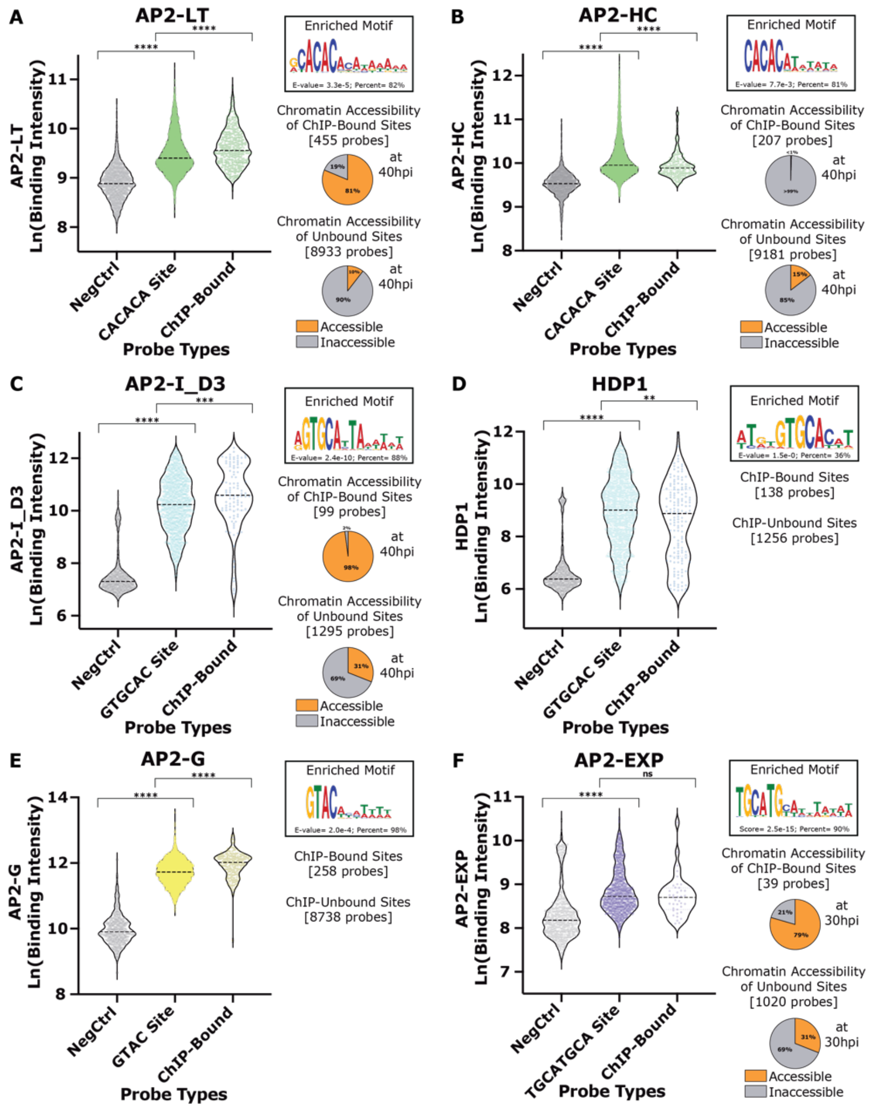


**<u>Supplemental Figure 15: Motif enrichment and chromatin accessibility of ChIP-bound and ChIP-unbound sites for six TFs</u>**

(**A**) *Left:* Binding intensity distributions for negative control sites (Grey), all CACACA motif sites (Green), and CACACA-containing sites that are AP2-LT ChIP-bound (Green). *Right Top:* AP2-LT ChIP-bound enriched motif, calculated E-value, and percent of occurrences. *Right Bottom:* Pie charts of chromatin accessibility and inaccessibility for ChIP-bound (top) and ChIP-unbound (bottom). Two-tailed Mann-Whitney test [p-value < 0.0001(****), 0.001(***), 0.01(**), or not significant(ns)]; (**B**) Binding intensity distributions, enriched motif and chromatin accessibility with AP2-HC data; (**C**) Binding intensity distributions, enriched motif and chromatin accessibility with AP2-I data; (**D**) Binding intensity distributions and enriched motif with HDP1 data. Chromatin accessibility data is not available for the life cycle stage HDP1 ChIP-seq was taken (stage II gametocytes); (**E**) Binding intensity distributions and enriched motif with AP2-G data. Chromatin accessibility data is not available for the life cycle stage AP2-G ChIP-seq was taken (committing gametocytes); (**F**) Binding intensity distributions, enriched motif and chromatin accessibility with AP2-EXP data.

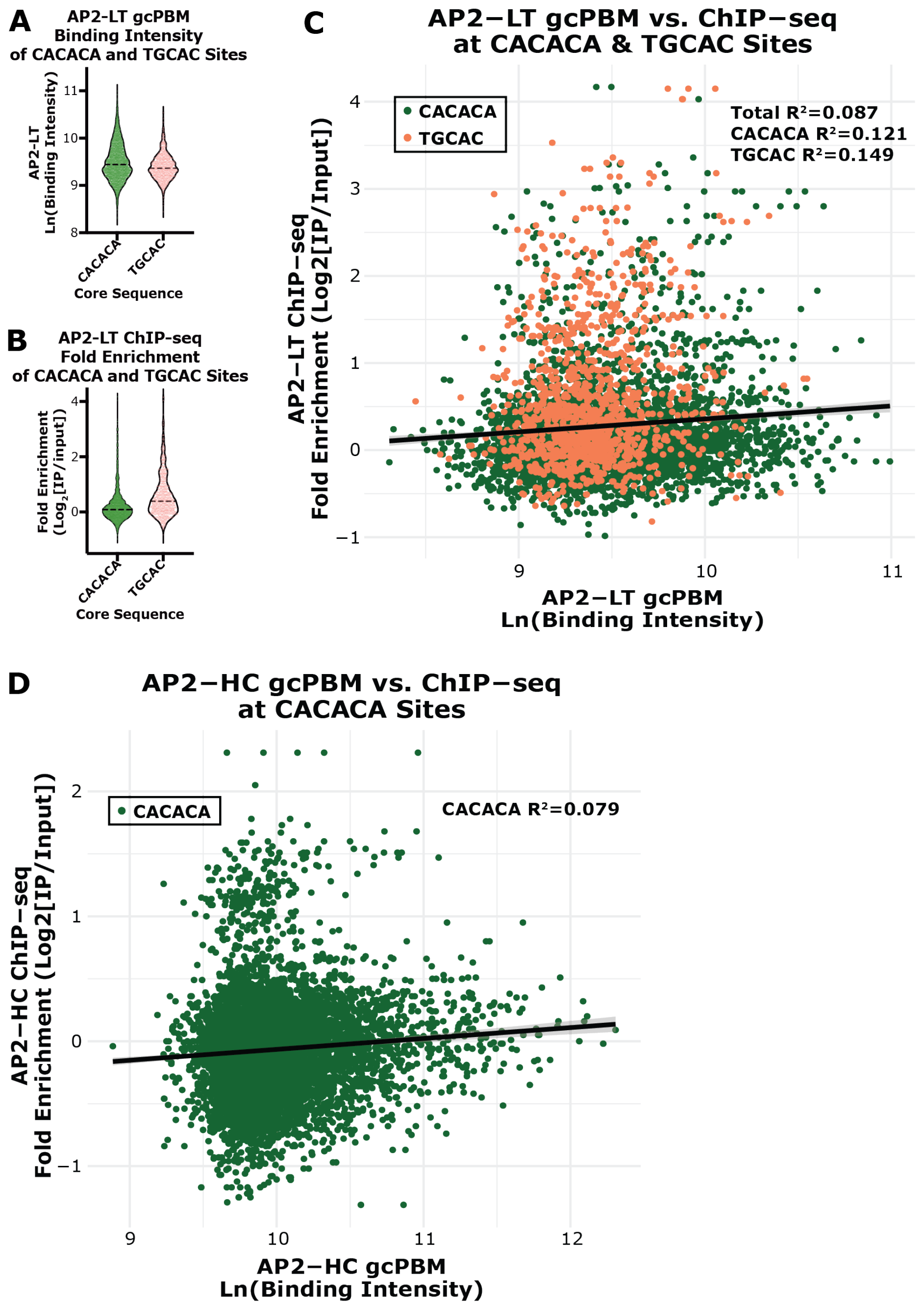


**<u>Supplemental Figure 16: Comparing AP2-LT and AP2-HC binding *in vitro* and *in vivo*</u>**

(**A**) AP2-LT gcPBM binding intensity across CACACA sites (Green) and TGCAC (Peach); (**B**) AP2-LT ChIP-seq fold enrichment ($Log_2$[IP/Input]) across CACACA sites (Green) and TGCAC (Peach); (**C**) Comparison of AP2-LT gcPBM binding intensity and AP2-LT ChIP-seq fold enrichment ($Log_2$[IP/Input]) across CACACA sites (Green) and TGCAC (Peach). *Top Right:* Pearson correlation values for all data points, only CACACA data points, and TGCAC data points. (**D**) Comparison of AP2-HC gcPBM binding intensity and AP2-HC ChIP-seq fold enrichment ($Log_2$[IP/Input]) across CACACA sites (Green). *Top Right:* Pearson correlation values for all data points.

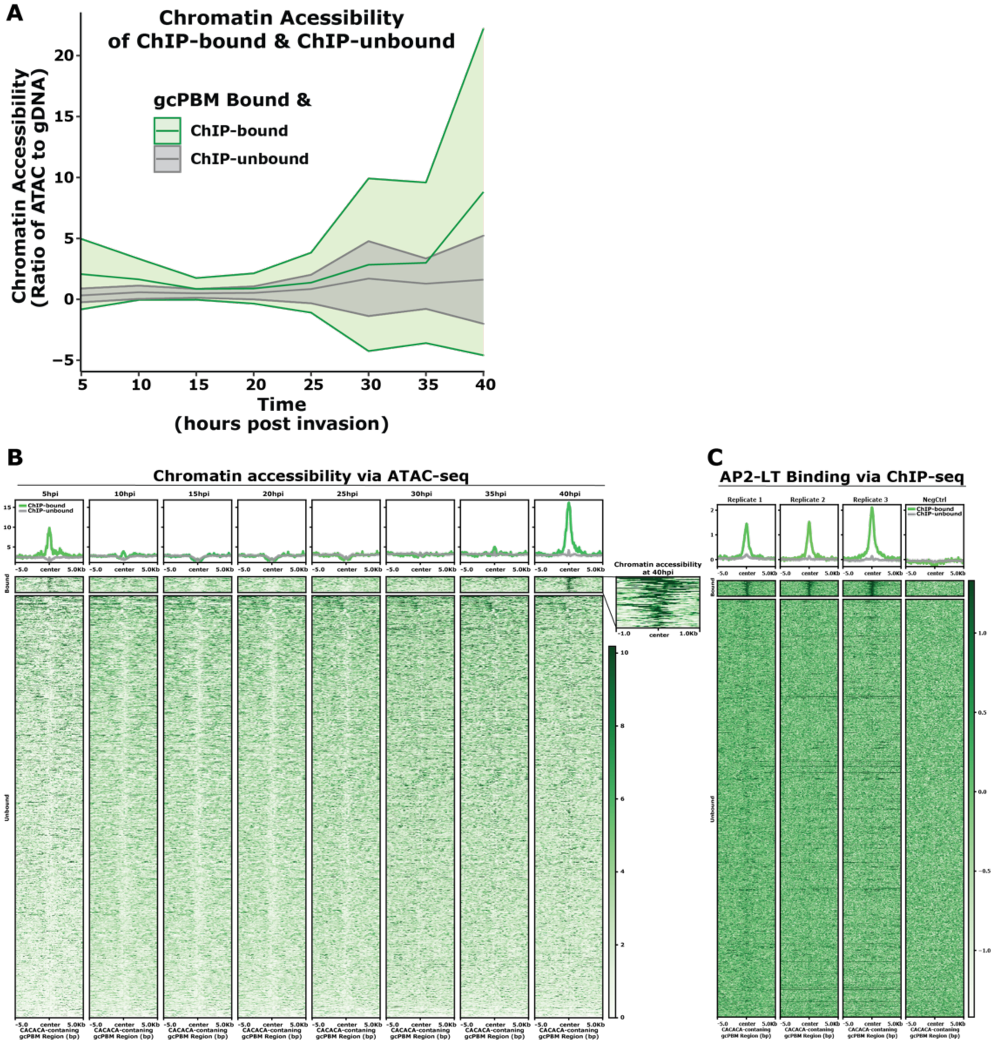


**<u>Supplemental Figure 17: Chromatin accessibility of AP2-LT ChIP-bound and ChIP-unbound CACACA-containing sites</u>**

(**A**) Line plot of chromatin accessibility across eight asexual stage timepoints (5hpi, 10hpi, 15hpi, 20hpi, 25hpi, 30hpi, 35hpi, and 40hpi) for AP2-LT ChIP-bound (Green) and ChIP-unbound (Grey) sites. Central line plotted is the median normalized read count over gDNA control. Upper and lower lines are the median plus and minus one standard deviation; (**B**) *Top:* Profile plot of the mean chromatin accessibility (median normalized read count over gDNA control) across eight asexual stage timepoints (5hpi, 10hpi, 15hpi, 20hpi, 25hpi, 30hpi, 35hpi, and 40hpi) for AP2-LT ChIP-bound (Green) and ChIP-unbound (Grey) sites. *Bottom:* Heatmap of chromatin accessibility (median normalized read count over gDNA control) across eight asexual stage timepoints (5hpi, 10hpi, 15hpi, 20hpi, 25hpi, 30hpi, 35hpi, and 40hpi) for AP2-LT ChIP-bound (top) and ChIP-unbound (bottom) sites. *Top Right zoom in:* Chromatin accessibility of AP2-LT ChIP-bound CACACA-containing sites at 40hpi; and (**C**) *Top:* Profile plot of the mean AP2-LT ChIP-seq fold enrichment ($Log_2$[IP/Input]) for all three biological replicates and no-epitope negative control sample across AP2-LT ChIP-bound (Green) and ChIP-unbound (Grey) sites. *Bottom:* Heatmap of the AP2-LT ChIP-seq fold enrichment ($Log_2$[IP/Input]) for all three biological replicates and no-epitope negative control sample AP2-LT ChIP-bound (Green) and ChIP-unbound (Grey) sites.

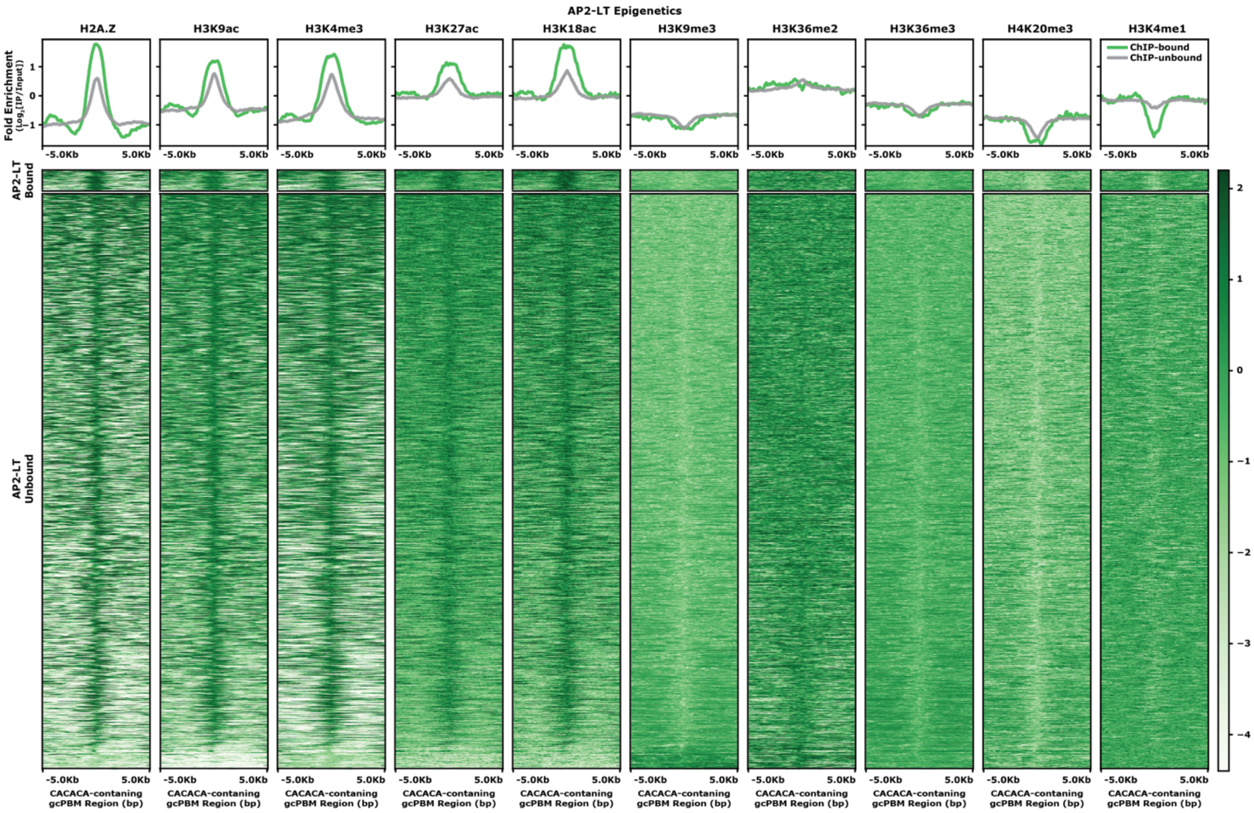


**Supplemental Figure 18: Epigenetic landscape of AP2-LT ChIP-bound and ChIP-unbound CACACA-containing sites**

*Top:* Profile plot of the mean ChIP-seq fold enrichment ($Log_2$[IP/Input]) of five activation epigenetic marks (H2A.Z, H3K9ac, H3K4me3, H3K27ac, and H3K18ac) and five repression epigenetic marks (H3K9me3, H3K36me2/3, H4K20me3, and H3K4me1) for AP2-LT ChIP-bound (Green) and ChIP-unbound (Grey) sites. *Bottom:* Heatmap of ChIP-seq fold enrichment ($Log_2$[IP/Input]) of five activation epigenetic marks (H2A.Z, H3K9ac, H3K4me3, H3K27ac, and H3K18ac) and five repression epigenetic marks (H3K9me3, H3K36me2/3, H4K20me3, and H3K4me1) for AP2-LT ChIP-bound (Green) and ChIP-unbound (Grey) sites.

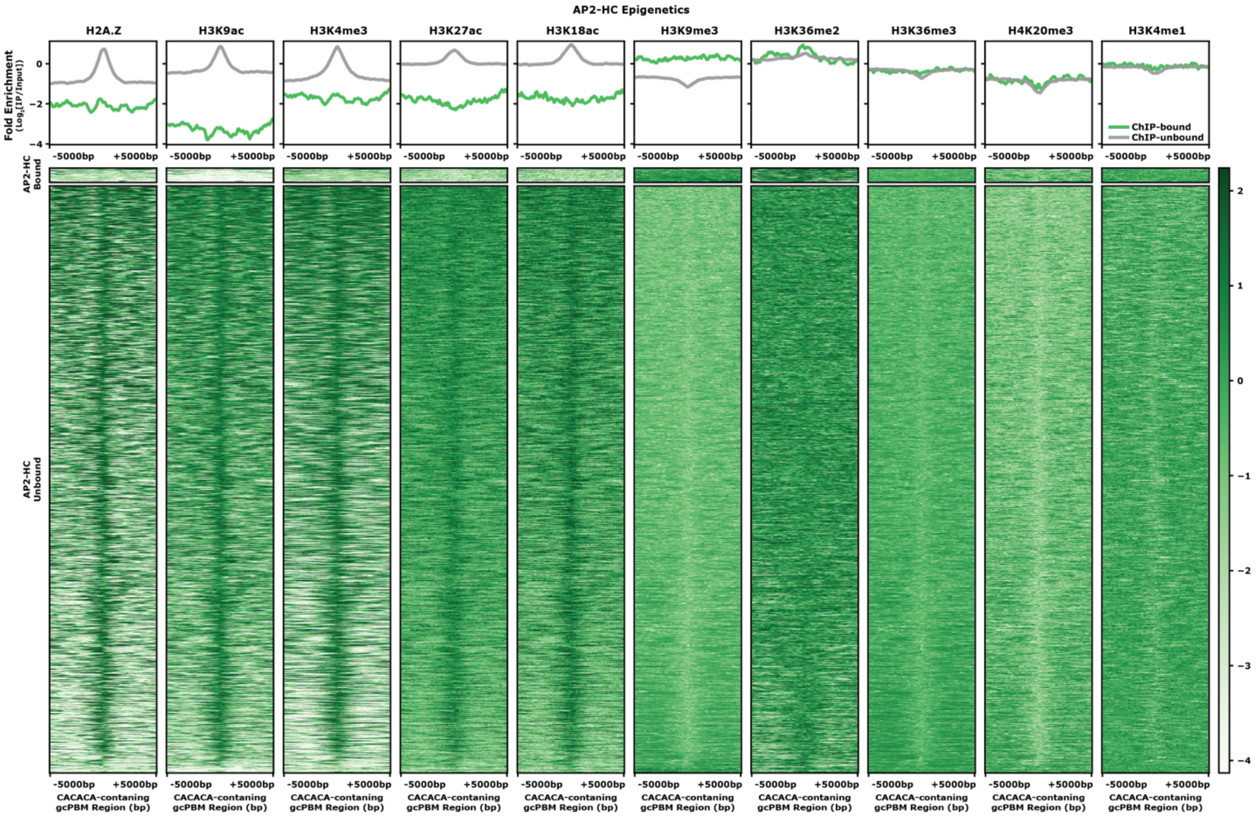


**Supplemental Figure 19: Epigenetic landscape of AP2-HC ChIP-bound and ChIP-unbound CACACA-containing sites**

*Top:* Profile plot of the mean ChIP-seq fold enrichment ($Log_2$[IP/Input]) of five activation epigenetic marks (H2A.Z, H3K9ac, H3K4me3, H3K27ac, and H3K18ac) and five repression epigenetic marks (H3K9me3, H3K36me2/3, H4K20me3, and H3K4me1) for AP2-HC ChIP-bound (Green) and ChIP-unbound (Grey) sites. *Bottom:* Heatmap of ChIP-seq fold enrichment ($Log_2$[IP/Input]) of five activation epigenetic marks (H2A.Z, H3K9ac, H3K4me3, H3K27ac, and H3K18ac) and five repression epigenetic marks (H3K9me3, H3K36me2/3, H4K20me3, and H3K4me1) for AP2-HC ChIP-bound (Green) and ChIP-unbound (Grey) sites.

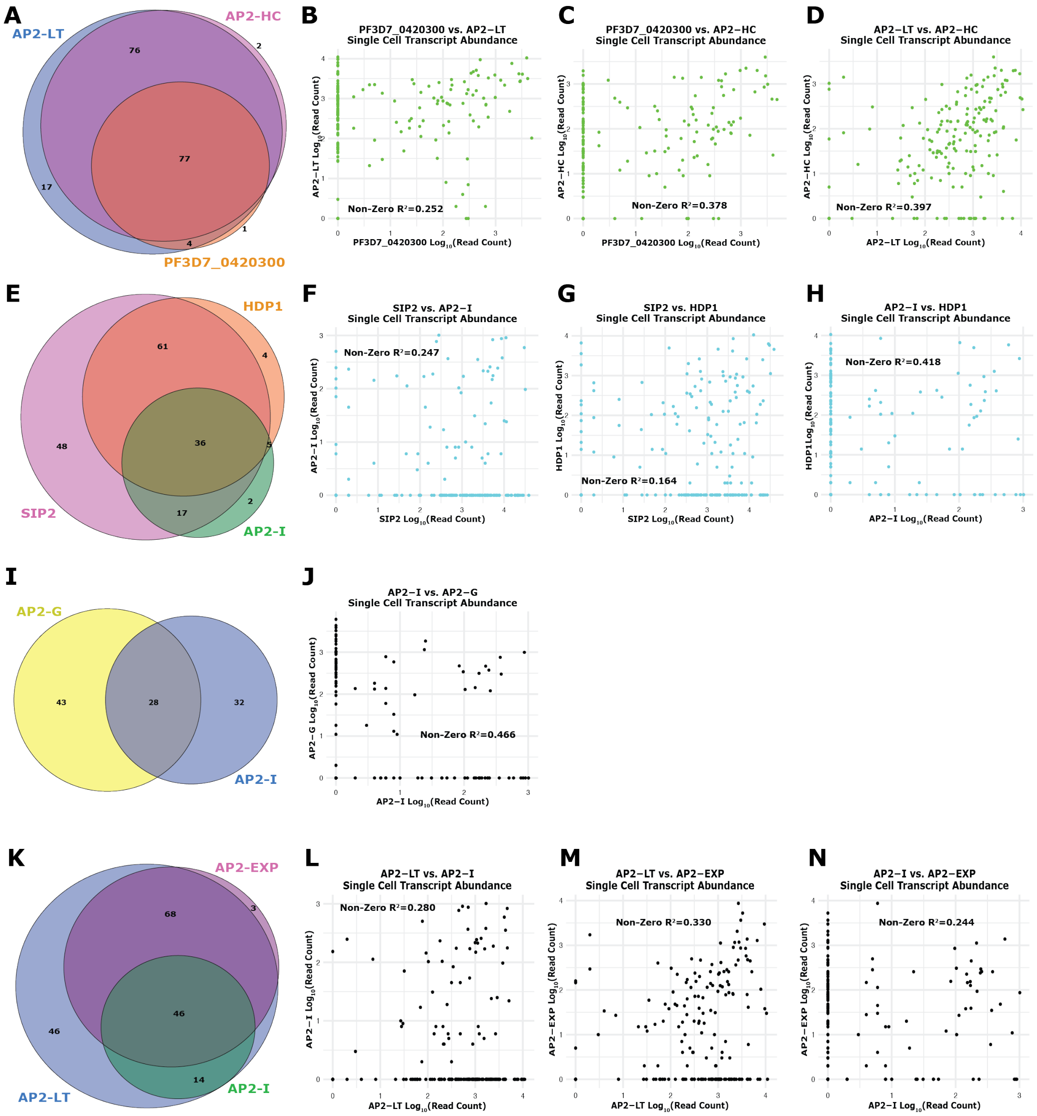


**Supplemental Figure 20: Single-cell transcript abundance data for all TF pairwise comparisons**

(**A**) Quantitative overlap of the number of single cells with each CACACA-binding TF transcribed (PF3D7_0420300, AP2-LT, and AP2-HC); (**B**) Comparison of $Log_{10}$(Read Count) for PF3D7_0420300 and AP2-LT. Pearson correlation of plotted data does not include zero values; (**C**) Comparison of $Log_{10}$(Read Count) for PF3D7_0420300 and AP2-HC; (**D**) Comparison of $Log_{10}$(Read Count) for AP2-LT and AP2-HC; (**E**) Quantitative overlap of the number of single cells with each GTGCAC-binding TF transcribed (SIP2, AP2-I, and HDP1); (**F**) Comparison of $Log_{10}$(Read Count) for SIP2 and AP2-I; (**G**) Comparison of $Log_{10}$(Read Count) for SIP2 and HDP1; (**H**) Comparison of $Log_{10}$(Read Count) for AP2-I and HDP1; (**I**) Quantitative overlap of the number of single cells with AP2-G and AP2-I transcribed; (**J**) Comparison of $Log_{10}$(Read Count) for AP2-I and AP2-G; (**K**) Quantitative overlap of the number of single cells with AP2-LT, AP2-I, and AP2-EXP; (**L**) Comparison of $Log_{10}$(Read Count) for AP2-LT and AP2-I; (**M**) Comparison of $Log_{10}$(Read Count) for AP2-LT and AP2-EXP; and (**N**) Comparison of $Log_{10}$(Read Count) for AP2-I and AP2-EXP.

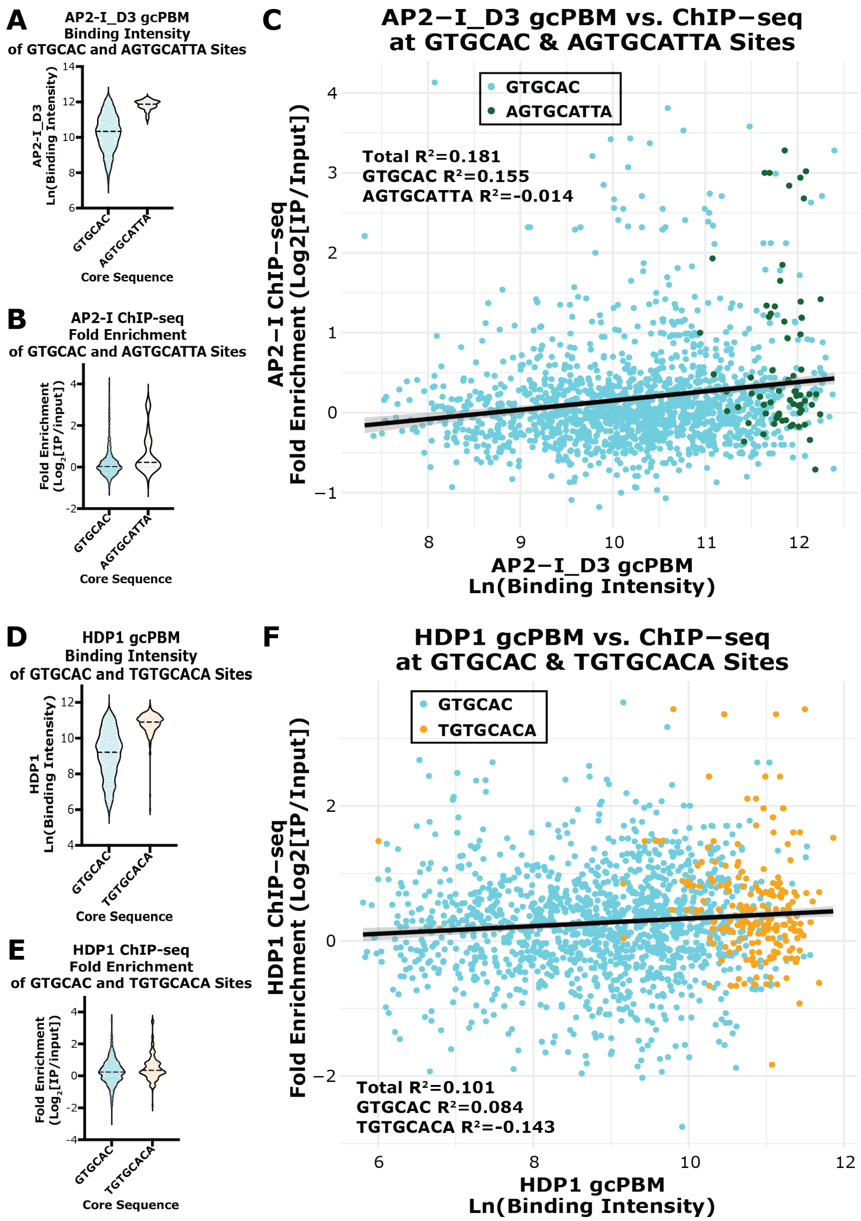


**<u>Supplemental Figure 21: Comparing AP2-I and HDP1 binding *in vitro* and *in vivo*</u>**

**(A)** AP2-I_D3 gcPBM binding intensity across GTGCAC sites (Blue) and AGTGCATTA (Green); **(B)** AP2-I ChIP-seq fold enrichment ($Log_2$[IP/Input]) across GTGCAC sites (Blue) and AGTGCATTA (Green); **(C)** Comparison of AP2-I_D3 gcPBM binding intensity and AP2-I ChIP-seq fold enrichment ($Log_2$[IP/Input]) across GTGCAC sites (Blue) and AGTGCATTA (Green). *Top Right:* Pearson correlation values for all data points, only GTGCAC data points, and AGTGCATTA data points; **(D)** HDP1 gcPBM binding intensity across GTGCAC sites (Blue) and TGTGCACA (Orange); **(E)** HDP1 ChIP-seq fold enrichment ($Log_2$[IP/Input]) across GTGCAC sites (Blue) and TGTGCACA (Orange); **(F)** Comparison of HDP1 gcPBM binding intensity and HDP1 ChIP-seq fold enrichment ($Log_2$[IP/Input]) across GTGCAC sites (Blue) and TGTGCACA (Orange). *Top Right:* Pearson correlation values for all data points, only GTGCAC data points, and TGTGCACA data points.

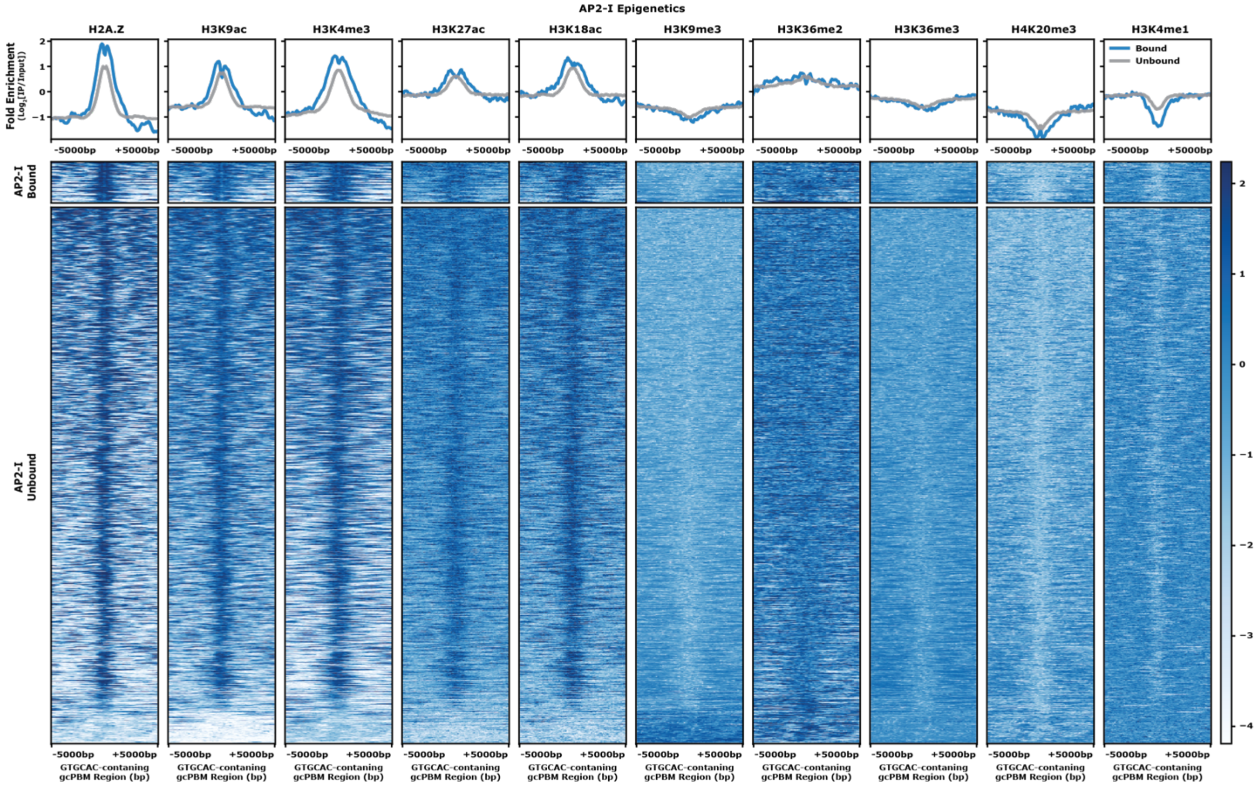


**Supplemental Figure 22: Epigenetic landscape of AP2-I ChIP-bound and ChIP-unbound GTGCAC-containing sites**

*Top:* Profile plot of the mean ChIP-seq fold enrichment ($Log_2$[IP/Input]) of five activation epigenetic marks (H2A.Z, H3K9ac, H3K4me3, H3K27ac, and H3K18ac) and five repression epigenetic marks (H3K9me3, H3K36me2/3, H4K20me3, and H3K4me1) for AP2-I ChIP-bound (Blue) and ChIP-unbound (Grey) sites. *Bottom:* Heatmap of ChIP-seq fold enrichment ($Log_2$[IP/Input]) of five activation epigenetic marks (H2A.Z, H3K9ac, H3K4me3, H3K27ac, and H3K18ac) and five repression epigenetic marks (H3K9me3, H3K36me2/3, H4K20me3, and H3K4me1) for AP2-I ChIP-bound (Blue) and ChIP-unbound (Grey) sites.

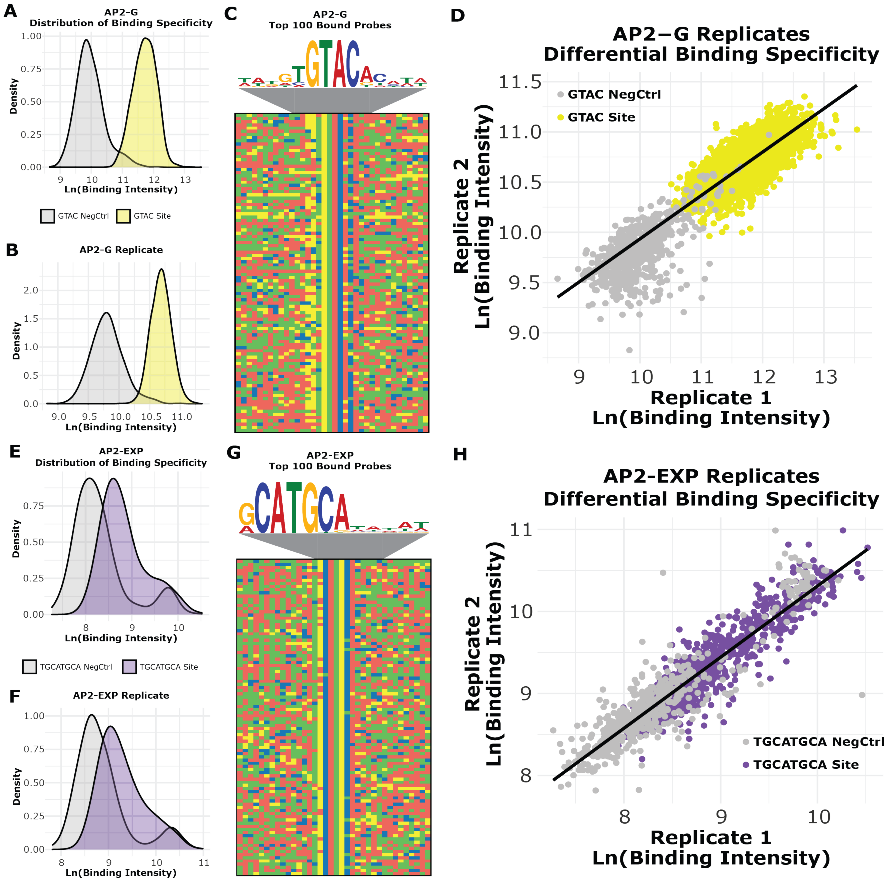


**<u>Supplemental Figure 23: Distribution of binding specificity and replicate correlation for AP2-G and AP2-EXP</u>**

(**A**) Density plot depicting the AP2-G binding intensities to the GTAC negative control sequences (Grey) and GTAC-containing sequences (Yellow); **(B)** Distributions of the binding intensities for AP2-G technical replicate (GTAC probes [Yellow] and negative control probes [Grey]); (**C**) The DNA motif enriched[86] in the top 100 GTAC probes bound by AP2-G with a four-color plot of the top 100 bound GTAC probes underneath. Color representations: A (Red), C (Blue), G (Yellow), and T (Green); (**D**) Scatter plot comparing the binding intensities for AP2-G technical replicates. Each data point represents the median binding intensity across all technical replicates of the TF to a specific DNA sequence: GTAC negative control sequences (Grey) and GTAC-containing sequences (Yellow); (**E**) Density plot depicting the AP2-EXP binding intensities to the TGCATGCA negative control sequences (Grey) and TGCATGCA-containing sequences (Purple); (**F**) Density plot depicting the AP2-EXP technical replicate binding intensities to the TGCATGCA negative control sequences (Grey) and TGCATGCA-containing sequences (Purple); (**G**) The DNA motif enriched[86] in the top 100 TGCATGCA probes bound by AP2-EXP with a four-color plot of the top 100 bound TGCATGCA probes underneath; and (**H**) Scatter plot comparing the binding intensities for AP2-EXP technical replicates. Each data point represents the median binding intensity across all technical replicates of the TF to a specific DNA sequence: TGCATGCA negative control sequences (Grey) and TGCATGCA-containing sequences (Purple); **(H)** *Left*: Distributions of the binding intensities for AP2-EXP technical replicate (TGCATGCA probes [Purple] and negative control probes [Grey]).

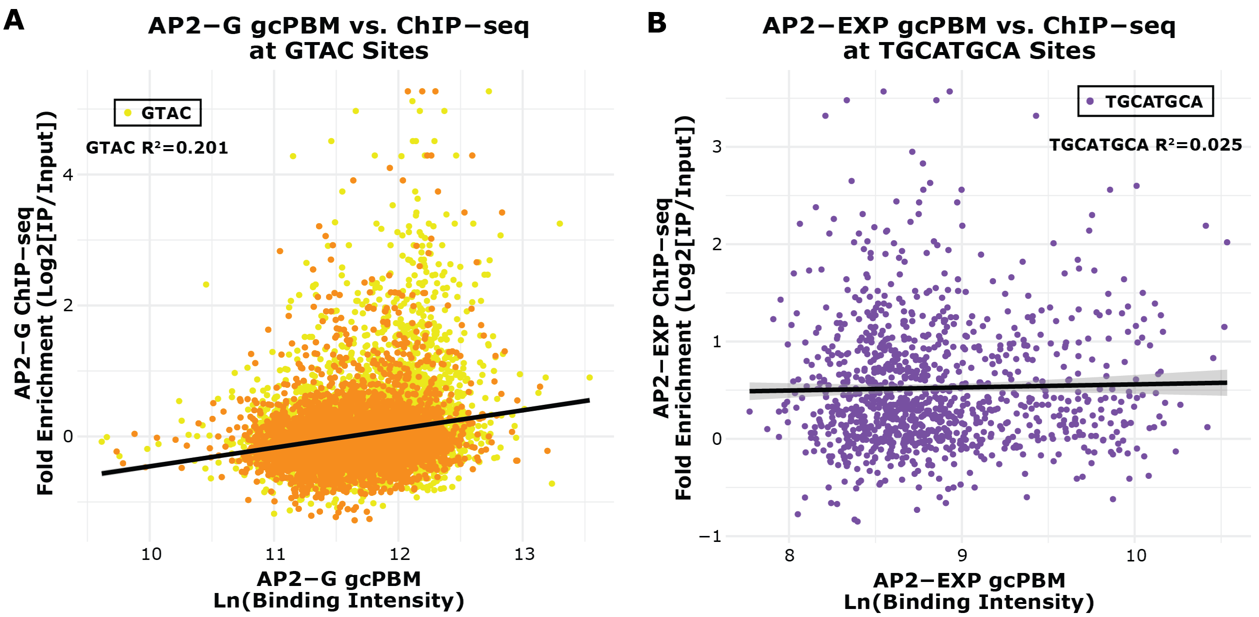


**Supplemental Figure 24: Comparing AP2-G and AP2-EXP binding *in vitro* and *in vivo***

**(A)** Comparison of AP2-G gcPBM binding intensity and AP2-G ChIP-seq fold enrichment ($Log_2$[IP/Input]) across GTAC sites (Yellow). *Top Right:* Pearson correlation values for all data points; **(D)** Comparison of AP2-EXP gcPBM binding intensity and AP2-EXP ChIP-seq fold enrichment ($Log_2$[IP/Input]) across TGCATGCA sites (Purple). *Top Right:* Pearson correlation values for all data points.

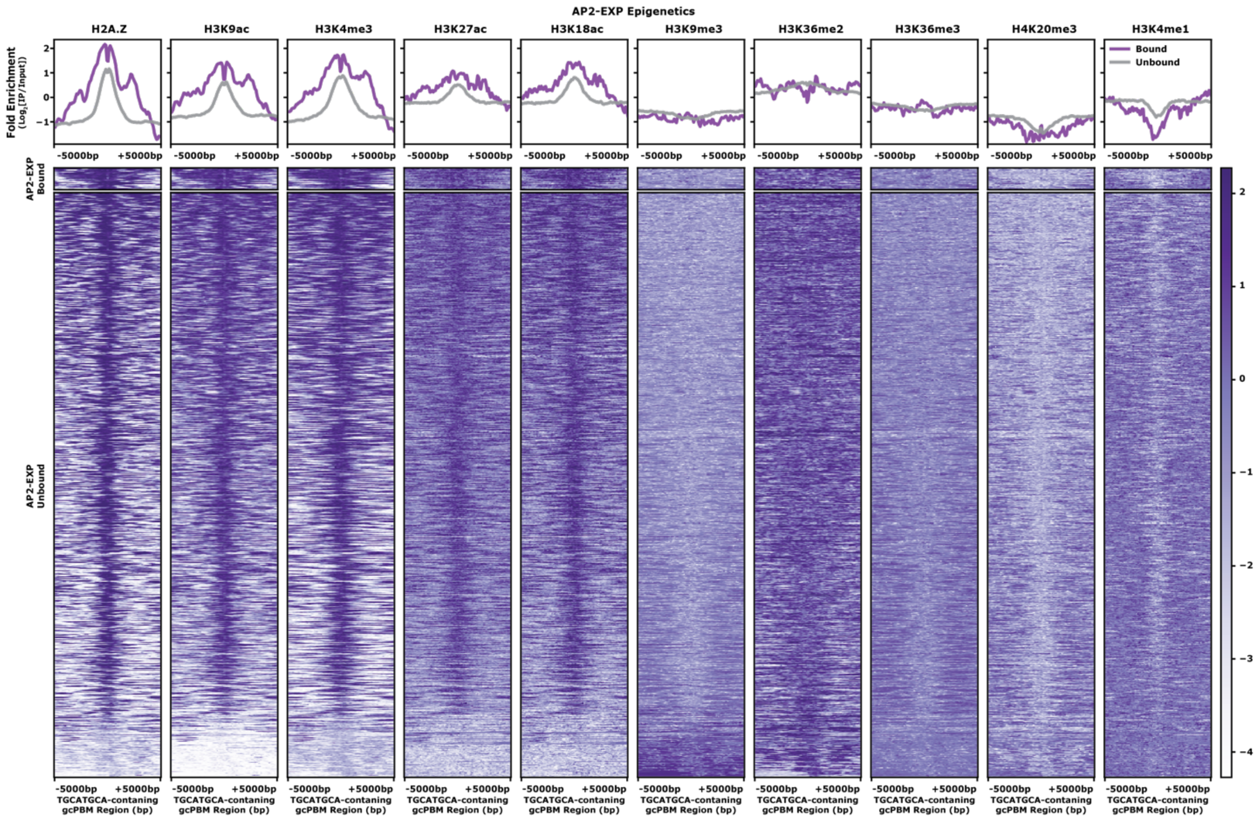


**Supplemental Figure 25: Epigenetic landscape of AP2-EXP ChIP-bound and ChIP-unbound TGCATGCA-containing sites**

*Top:* Profile plot of the mean ChIP-seq fold enrichment ($Log_2$[IP/Input]) of five activation epigenetic marks (H2A.Z, H3K9ac, H3K4me3, H3K27ac, and H3K18ac) and five repression epigenetic marks (H3K9me3, H3K36me2/3, H4K20me3, and H3K4me1) for AP2-EXP ChIP-bound (Purple) and ChIP-unbound (Grey) sites. *Bottom:* Heatmap of ChIP-seq fold enrichment ($Log_2$[IP/Input]) of five activation epigenetic marks (H2A.Z, H3K9ac, H3K4me3, H3K27ac, and H3K18ac) and five repression epigenetic marks (H3K9me3, H3K36me2/3, H4K20me3, and H3K4me1) for AP2-EXP ChIP-bound (Purple) and ChIP-unbound (Grey) sites.

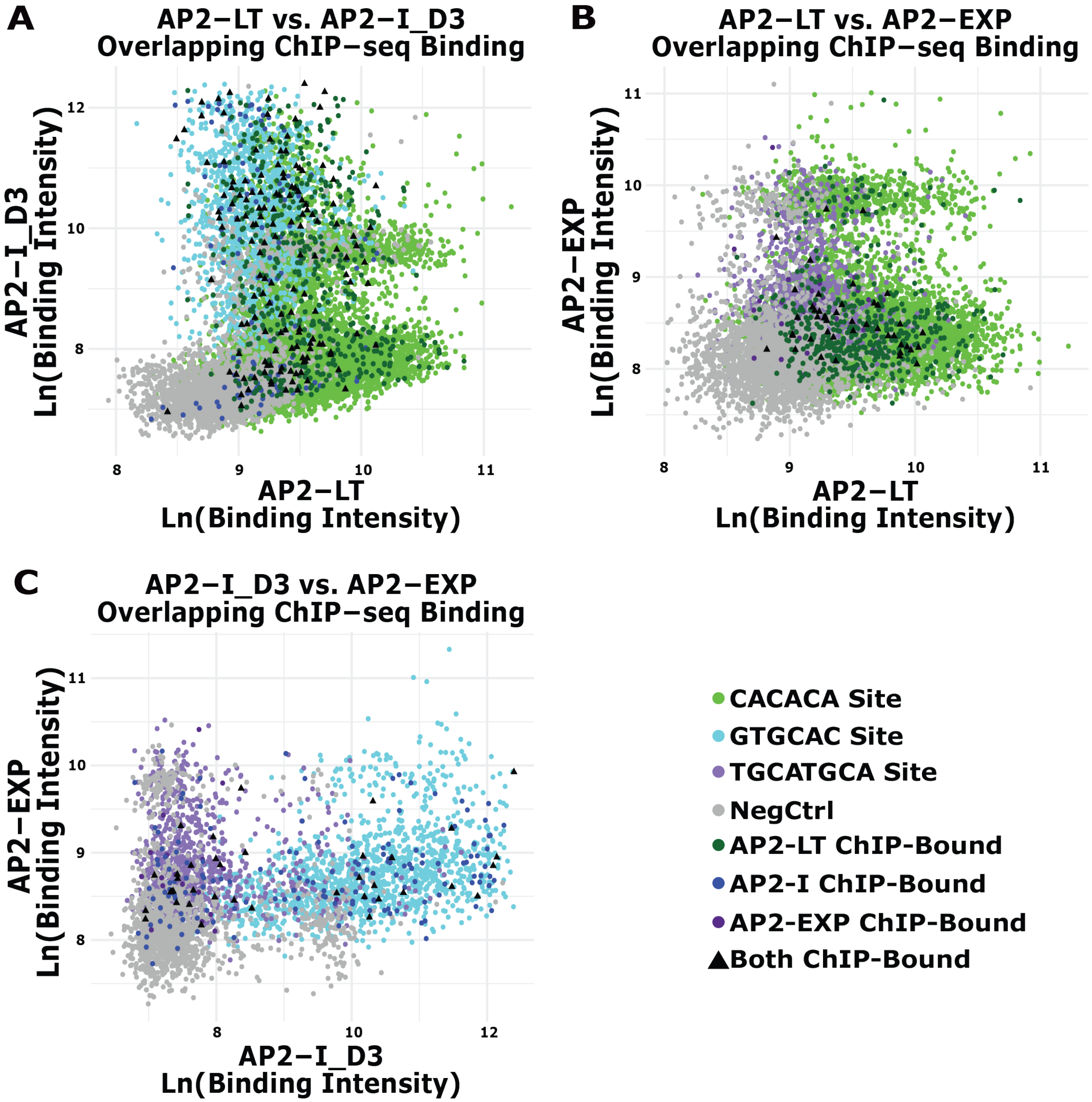


**Supplemental Figure 26: Overlapping *in vitro* binding preferences across DNA motif types**
(**A**) Comparison of the gcPBM binding intensities for AP2-LT and AP2-I_D3. Negative control sequences (Grey), CACACA-containing sequences (Green), GTGCAC-containing sequences (Blue), TGCATGCA-containing sequences (Purple), AP2-LT ChIP-bound sequences (Dark Green), AP2-I ChIP-bound sequences (Dark Blue), AP2-EXP ChIP-bound sequences (Dark Purple), and AP2-LT/AP2-I, AP2-LT/AP2-EXP, or AP2-I/AP2-EXP co-bound ChIP-bound sequences (Black triangle); (**B**) Comparison of the binding intensities for AP2-LT and AP2-EXP; and (**C**) Comparison of the binding intensities for AP2-I_D3 and AP2-EXP.
